# Karrikin and strigolactone signalling affect pattern-triggered immunity and resistance to specific pathogens

**DOI:** 10.64898/2026.05.05.722863

**Authors:** Sebastian D. Schade, Kees Buhrman, Anne Schneider, Debangana Moitra, Athanasios Makris, Hua Wei, Rosa Lozano-Durán, Martina K. Ried-Lasi, Birgit Kemmerling, Caroline Gutjahr, Martin Stegmann

## Abstract

Hormone signalling is important for plant adaptation to biotic stress. Karrikins (KARs), smoke-derived compounds, and strigolactones (SLs), endogenous plant hormones, are families of butenolide molecules, sharing a convergent perception and signalling pathway to regulate a plethora of developmental processes and plant-symbiont relationships. Perception of KARs and SLs is mediated by the α/β-hydro-lase KARRIKIN INSENSITIVE 2 (KAI2) and DWARF14 (D14), respectively, each resulting in the formation of an E3 ubiquitin ligase complex with the F-Box protein MORE AXILLIARY GROWTH 2 (MAX2) to target transcriptional repressors of the SUPPRESSOR OF MAX2 (SMAX)/SMAX-LIKE (SMXL) family for degradation. Most likely, KAI2 additionally perceives a still elusive endogenous ligand (KAI2-ligand, KL). Recent reports suggest a role of KL/SL signalling in plant immunity, but how these pathways are involved in defence, while balancing appropriate symbiont interactions remains largely unknown. Here, we report that KL and SL signalling quantitatively modulate plant immune responses and pathogen resistance. In *Arabidopsis thaliana* (hereafter Arabidopsis), disrupting or de-repressing KL or SL signalling affects plant susceptibility to a variety of plant pathogens. Furthermore, we describe a previously unknown role for KL and SL signalling in modulating pattern-triggered immunity (PTI). Interfering with KL and SL perception in Arabidopsis had similar effects on microbe-associated molecular pattern (MAMP)-triggered reactive oxygen species production, but transcriptomic profiling suggests a predominant role for KL signalling in regulating the extent of PTI. Importantly, KAI2- and D14-mediated regulation of MAMP-triggered ROS production extends to *Lotus japonicus* and, in the case of KAI2, to *Nicotiana benthamiana*, indicating conserved immuno-modulatory roles across dicotyledonous lineages. Together our data identify KL and SL signalling, with a predominant role for the KL pathway, as a conserved modulatory layer of plant immunity and provides a framework for understanding how developmental pathways intersect with immune regulation.

## Introduction

Plants are continuously exposed to a wide range of environmental challenges of both abiotic and biotic origin, which can adversely affect plant growth, development, yield and survival (1). Consequently, plants have evolved sophisticated and flexible adaptive strategies encompassing morphological, metabolic and transcriptional responses. Many of these mechanisms are regulated by phytohormones, tightly coordinating the balance between plant developmental processes and stress adaptation. In the context of biotic stress, several phytohormones have been identified as central regulators of plant defence, with salicylic acid (SA), jasmonic acid (JA), and abscisic acid (ABA) playing a predominant role (2). Additional hormones such as ethylene, brassinosteroids, auxins, and gibberellins, further fine-tune plant immune responses by contributing to the complex crosstalk among defence signalling networks (3). In recent years, the karrikin (KAR) and strigolactone (SL) pathways have emerged as potential hormonal pathways sharing similar cross-regulatory functions (4).

KARs are exogenous butenolides present in the smoke of burnt vegetation that promote seed germination and early seedling growth after wildfire events (5–7). Beyond these effects, the KAR signalling pathway is involved in enhancing seedling responses to light (8), modulating drought resistance (9,10), and regulating root and root hair development (11–13). Perception of KAR is mediated by the non-canonical ɑ/β-hydrolase KARRIKIN INSENSITIVE2 (KAI2), however, KAR likely requires metabolization to facilitate KAI2 binding (14,15). Genetic evidence suggests the existence of a yet unidentified plant endogenous hormone ligand for KAI2 (KAI2-ligand, KL), which likely accounts for the widespread presence of KAR/KL signalling pathways in plants from non-fire-prone environments (16,17).

Upon ligand binding, KAI2 interacts with MORE AXILLARY GROWTH2 (MAX2), an F-box component of a Skp-Cullin-F-box (SCF) E3 ubiquitin ligase complex (18), to recruit SUPPRESSOR OF MAX2 1 (SMAX1) and, in Arabidopsis, the partially redundant SMAX1-LIKE2 (SMXL2) (19,20). These proteins belong to the SMAX1 clade of the SMXL proteins (21), and their association stabilises the ligase-recep-tor-substrate complex (22). This dynamic interaction results in the polyubiquitination of SMAX1/SMXL2, followed by degradation via the 26S proteasome (23–25). Proteasome-mediated degradation of SMXL proteins upon hormone perception then initiates downstream signalling responses by relieving transcriptional repression.

SLs comprise a group of endogenous, carotenoid-derived butenolides that regulate diverse aspects of plant physiology and are widespread throughout the plant kingdom (26). SLs influence root architecture (27,28), repress shoot branching (29,30), promote leaf senescence (31), and positively regulate secondary growth (32). SL perception is mechanistically analogous to KAR signalling as it is mediated by DWARF14 (D14) (33,34), a paralogue of the more ancient KAI2 (18,35,36). Ligand-bound D14 similarly interacts with SCF^MAX2^ to target the redundant growth regulators SMXL6, SMXL7 and SMXL8, which form the SMXL68 clade of SMXL proteins (37,38). Members of this clade exhibit partially overlapping expression patterns with SMAX1 and can display antagonistic effects (37), underscoring the context-dependent regulation mediated by SMXL proteins. To confer functional specificity, members of the SMXL protein family have been shown to regulate distinct groups of genes through direct DNA binding and/or selective interactions with transcription factors and transcriptional co-repressors, including TOPLESS (TPL) and TOPLESS-RELATED (TPR) proteins (39–43).

Beyond their roles in growth regulation, both KAR and SL signalling pathways play important functions in plant-microbe interactions. KL perception is required for the establishment of symbiotic relationships with arbuscular mycorrhizal (AM) fungi in symbiosis-forming plant species (44–49). SLs in turn are actively secreted into the rhizosphere under nutrient-limiting conditions, where they promote the activity of AM fungi development and potentially provide a directional signal for root recognition (50–53).

Increasing evidence challenges the traditional view of symbiosis and defence as strictly separate processes, revealing extensive overlap between the pathways that distinguish beneficial symbionts from pathogens. Rather than acting in isolation, converging gene and signalling modules tightly cross-regulate plant-microbe interactions to balance symbiosis and immunity (54–60).

To detect and withstand pathogen infection, plants have evolved a multi-layered innate immune system (61). The primary layer of induced plant immunity is referred to as pattern-triggered immunity (PTI), that relies on plasma membrane localised pattern recognition receptors (PRRs) for detection of conserved microbe-associated molecular patterns (MAMPs) (62). Recognition of MAMPs activates signalling responses, including MITOGEN-ACTIVATED PROTEIN KINASE (MAPK) cascades, the production of reactive oxygen species (ROS), transcriptional reprogramming and stomatal closure, enabling plants to fend off most potential pathogens (63,64). Previous studies have suggested roles for KL and SL signalling in immune- and defence-related processes (65–67); however, the underlying mechanisms remain largely elusive.

Here, we investigated the role of KL and SL signalling pathways in modulating plant immunity across a variety of dicotyledonous plant lineages. We challenged Arabidopsis KL and SL signalling mutants with a diverse set of pathogens spanning multiple kingdoms. Our data reveal distinct pathogen-dependent involvement of KL signalling for infection outcomes on Arabidopsis, while SL signalling appears to play no role. However, we demonstrate that both KL and SL signalling mutants are defective in MAMP-triggered production of ROS. By RNA-sequencing (RNAseq) and MAMP-induced resistance experiments, we demonstrate a predominant role for KL signalling in regulating the extent of PTI. Finally, we show that compromised ROS production extends to *Lotus japonicus* mutants defective in KL and SL signalling and *kai2* mutants in *Nicotiana benthamiana*, suggesting a conserved regulatory module across diverse dicotyledonous species. Our data thus unravel a novel role for KL and SL signalling in the regulation of pathogenic plant-microbe-interactions and PTI with a predominant role for the KL pathway.

## Results

### Karrikin and strigolactone signalling affect resistance to specific pathogens

Previous reports demonstrated that both SL production or exogenous application of KARs and SLs can influence plant-pathogen interactions (67–69). To determine how KL and SL perception pathways affect host-pathogen interactions, we performed infection assays in Arabidopsis KL and SL signalling mutants, namely the KL and SL receptor mutants *kai2* and *d14*, the mutant of the KL/SL-shared F-box protein *max2* and mutants in the proteolytic targets of the two pathways *smax1 smxl2* and *smxl6,7,8* respectively, with pathogens spanning multiple kingdoms of life.

Infection of all genotypes with the obligate biotrophic powdery mildew fungus *Erysiphe cruciferarum* (*Ecr*), revealed that the KL receptor mutant *kai2-2* and the F-box protein mutant *max2-1* were more resistant, illustrated by the reduced numbers of conidiophores (*Ecr* reproductive structures) compared to the Col-0 control (Fig. 1A, B). Surprisingly, and in contrast to the canonical KL and SL signalling pathway configuration, the triple mutant of the proteolytic targets of SL signalling *smxl6,7,8* displayed a similarly increased resistance to *Ecr*, whereas neither the SL receptor *d14-1* mutant nor the mutant in canonical proteolytic targets of KL signalling *smax1 smxl2* displayed altered *Ecr* susceptibility. These findings suggest that KL signalling acts as a susceptibility factor during *Ecr* infection, potentially in a non-canonical manner.

**Figure 1:**
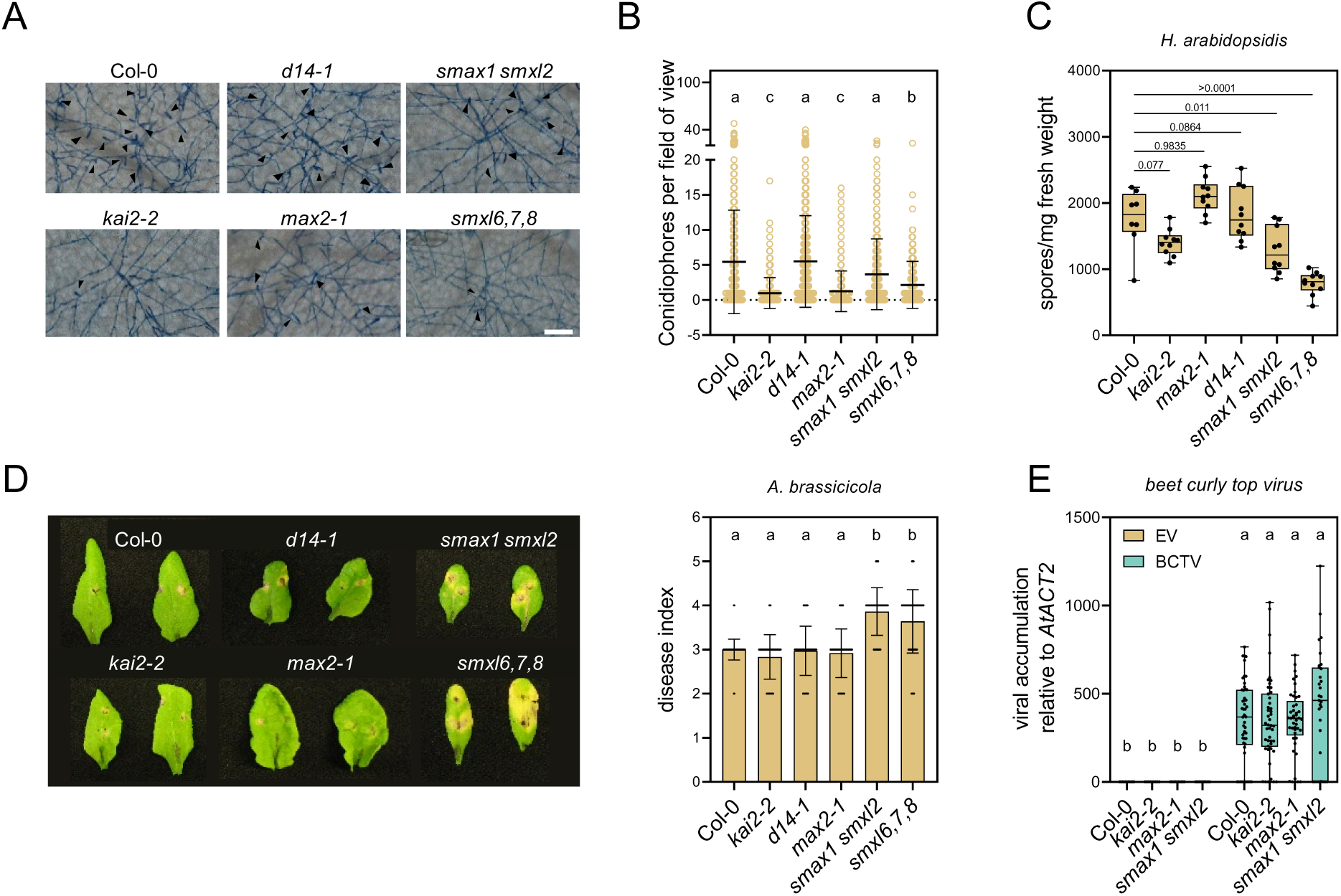
KL signalling modulates diverse plant-pathogen interactions. **A)** Representative ink-stained *Ecr* colonies 5 days post-infection (dpi) on mature plants of the indicated genotypes. Black ar-rows indicate conidiophores. Scale bar represents 100 µm. **B)** Number of conidiophores per field of view (5 dpi) of *Ecr* colonies grown upon infection of the indicated genotypes. Mean ± SD, n ≥ 249, pooled from four independent experiments (Kruskal-Wallis test; Dunn’s post-hoc test, a/c-b: p < 0.005; a-c: p < 0.0001). **C)** Number of spores of *H. arabidopsidis* counted on 12-day-old plants of the indicated geno-types per mg fresh weight 6 dpi. Mean ± SD, n ≥ 8, data from one experiment, which was performed two more times with similar results (one-way ANOVA, Dunnett’s post-hoc test; p-values are indicated in the figure). **D)** Representative images of detached leaves of the indicated Arabidopsis genotypes (left) and disease index (right) 13 dpi with *A. brassicicola*. Mean ± SD, n = 72, pooled from 3 independent experiments (one-way ANOVA, Tukey’s post-hoc test; a-b: p < 0.0001). **E)** Accumulation of viral DNA scored by qPCR in leaves of 2- to 3-week-old Arabidopsis plants infected with BCTV at 21 dpi. Mean ± SD, n = 9 - 31, pooled from 4 independent experiments (one-way ANOVA, Tukey’s post-hoc test; a-b: p < 0.0001).

To understand whether KL and/or SL signalling is involved in other obligate biotrophic pathogen interactions, we infected plants with the biotrophic oomycete *Hyaloperonospora arabidopsidis* (*Hpa*). In contrast with our results from *Ecr* infections, we observed no significantly increased resistance in *kai2* and *max2* mutants (Fig. 1C; S1A). However, both *smax1 smxl2* and *smxl6,7,8* were more resistant to *Hpa*. This raises the question whether SMXL proteins may contribute to resistance against oomycetes independently of canonical KAI2-MAX2 and D14-MAX2 signalling.

Infections with necrotrophic fungi produced distinct phenotypes. Upon infection with *Alternaria brassicicola* or *Botrytis cinerea, kai2-2, d14-1* and *max2-1* mutants did not exhibit altered disease phenotypes. However, *smxl6,7,8* displayed increased symptom development to both necrotrophic fungi (Fig. 1D; S1B). In addition, *smax1 smxl2* displayed increased susceptibility upon *A. brassicicola* compared to the wild type (Fig. 1D), but not upon *B. cinerea* infection (Fig. S1B). This further supports the existence of KAI2-MAX2 and D14-MAX2 independent signalling through SMXL proteins.

Given that the observed infection phenotypes largely correlated with impaired KL signalling, we examined KL pathway mutants for altered susceptibility to the single-stranded DNA beet curly top virus (BCTV) (family *Geminiviridae*) (Fig. 1E). However, neither of the KL signalling mutants displayed altered infection phenotypes. Collectively, our data indicate pathogen specific effects of KL or SL signalling on susceptibility against diverse pathogens.

### Karrikin and strigolactone signalling pathways influence pattern-triggered immunity responses

Primary pathogen perception is mediated by PTI and its regulation by plant hormones, such as SA and JA, shapes infection outcomes (2). We therefore addressed the question whether KL and SL signalling may be involved in PTI.

We quantified ROS production after elicitation with the MAMP flg22, a conserved 22-amino-acid epitope derived from bacterial flagellin (70). Compared to Col-0, *kai2-2*, *d14-1* and *max2-1* displayed significantly reduced ROS accumulation (Fig. 2A, B). We observed comparable reduction in ROS production responses after treatment of plants with a diverse set of MAMPs, including bacterial elf18 (derived from elongation factor Tu), fungal chitin or the broadly conserved nlp20 peptide (derived from NECROSIS AND ETHYLENE-INDUCING PEPTIDE1-like proteins) (Fig. 2C). This suggests a general impairment of

**Figure 2:**
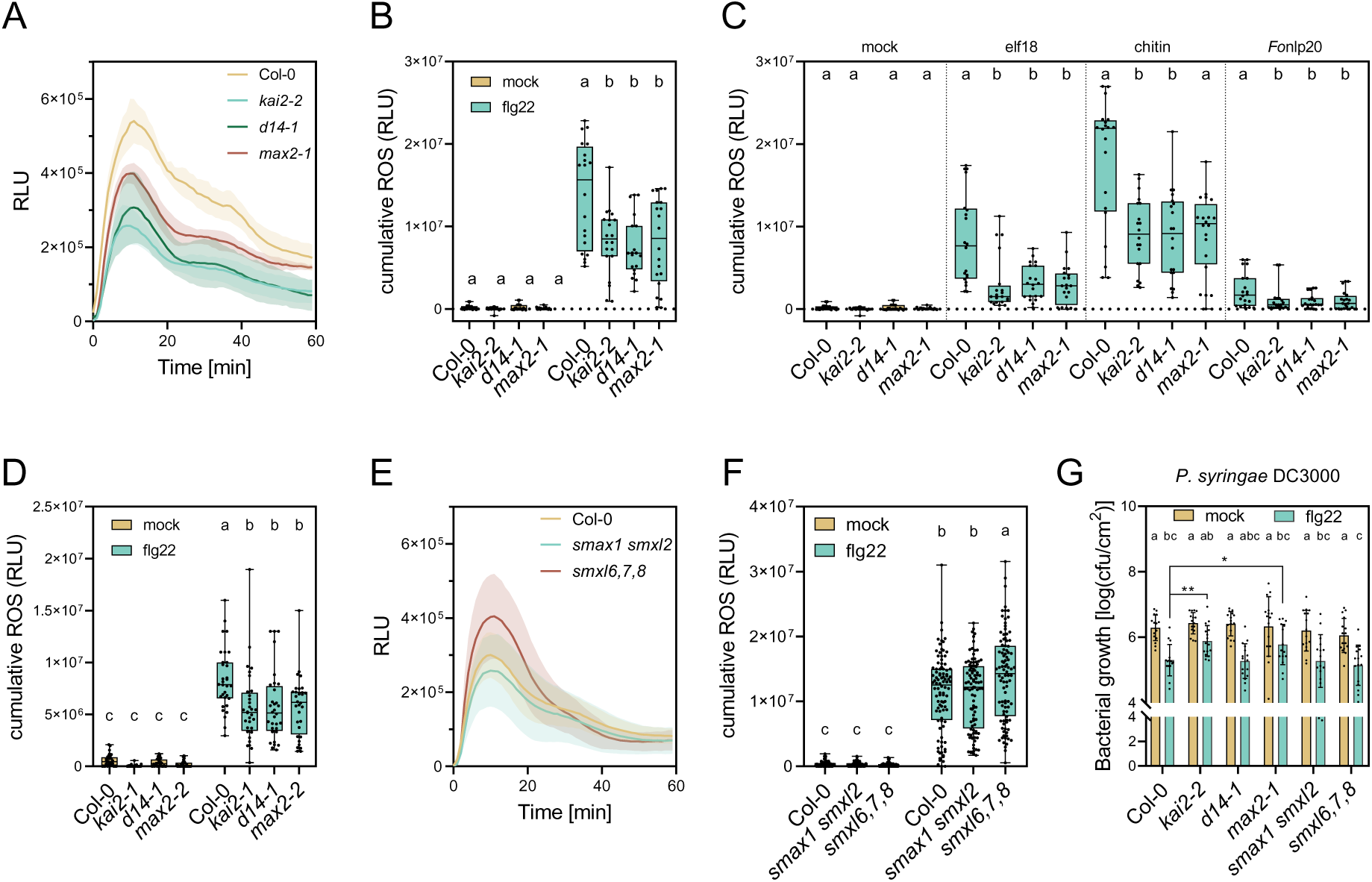
KL and SL signalling components modulate distinct PTI responses. A-F) Production of reactive oxygen species (ROS) in leaf discs of the indicated genotypes represented in relative light units (RLU). **A)** Accumulation kinetics after elicitation with flg22 (100 nM) over 60 min; mean ± SD, n = 4. **B)** Total RLU after elicitation with flg22 (100 nM); mean ± SD, n = 20, pooled from 5 independent replicates (one-way ANOVA, Tukey’s post-hoc test; a-b: p ≤ 0.002; a/b-c: p < 0.0001). **C)** Total RLU after elicitation with flg22 (100 nM); mean ± SD, n = 32, pooled from four independent replicates (one-way ANOVA, Tukey’s post-hoc test; a-b-c, p < 0.0001). **D)** Total RLU after treatment with elf18 (100 nM), chitin (20 ng/µl), Pep1 (1 µM) or *Fo*nlp20 (1 µM). Mean ± SD, n = 20, pooled from five biological replicates (one-way ANOVA, Tukey’s post-hoc test, performed separately for each treatment, a-b: p ≤ 0.045). **E)** ROS kinetics after elicitation with flg22 (100 nM); mean ± SD, n = 16. **F)** Total RLU after elicitation with flg22 (100 nM); mean ± SD, n = 96, with data pooled from six independent replicates (one-way ANOVA, Tukey’s post-hoc test; a-b-c: p < 0.002). **G)** flg22 induced resistance against *Pto* DC3000 infection. Leaves of the indicated genotypes were infiltrated with mock or flg22 (1 µM) before syringe infiltration with *Pto* DC3000 after 24 hours. 2 days post-inoculation (2 dpi) leaf discs were sampled to determine bacterial titres by counting colony forming units (CFU); Mean ± SD, n = 14-15, data pooled from four independent replicates (two-way ANOVA, Tukey’s post-hoc test, a-b-c: p < 0.02; two-tailed Student’s t-test, *: p < 0.03, **: p < 0.002).

MAMP-triggered ROS bursts in KL and SL perception mutants. Reduced ROS production upon flg22 treatment in *kai2* and *max2* mutants was confirmed with the independent alleles *kai2-1* and *max2-2* (Fig. 2D). In contrast, *smax1 smxl2* showed unaltered, while *smxl6,7,8* had significantly enhanced flg22-triggered ROS production (Fig. 2E, F), indicating specific regulatory roles for distinct SMXL clades in controlling PTI-associated ROS production.

To determine whether altered ROS production resulted from impaired protein accumulation or stability of key PTI signalling proteins, we analysed accumulation of the flg22 receptor FLAGELLIN SENSITIVE2 (FLS2) (71) and the chitin binding CHITIN ELICITOR RECEPTOR KINASE1 (CERK1) (72) by immunoblotting. Neither FLS2 nor CERK1 levels were changed in KL/SL signalling mutants (Fig. S2A, C). We also analysed levels of the PTI-associated ROS producing NADPH oxidase RESPIRATORY BURST OXIDASE HOMOLOG PROTEIN D (RBOHD) (73). RBOHD levels displayed a trend towards reduced accumulation across KL and SL mutants. However, these changes did not correlate with the observed ROS phenotypes (Fig. S2B, C), as *max2-1*, *d14-1*, *smax1 smxl2* and *smxl6,7,8* mutants all displayed reduced RBOHD levels despite exhibiting opposing MAMP-triggered ROS responses.

We next assessed MAPK activation upon flg22 treatment to test whether other early PTI responses are affected in SL and/or KL signalling mutants. However, we detected no differences in MAPK phosphorylation between genotypes (Fig. S3A, B), raising the question whether PTI modulation by KL and SL signalling is output specific.

Another hallmark of PTI is the ability to mount MAMP-induced resistance (74). Prior MAMP exposure establishes a primed defensive state that enhances resistance against subsequent pathogens, such as the bacterial pathogen *Pseudomonas syringae*. While growth of *P. syringae pv. tomato* (*Pto*) strain DC3000 did not differ between mutants and wild type under mock conditions, flg22 pre-treatment showed reduced induced resistance in *kai2-2* and *max2-1* compared to the other genotypes (Fig. 2G). Interestingly, despite exhibiting ROS burst phenotypes that resembled those of *kai2* and *max2-1*, *d14-1* did not display defects in flg22-induced resistance. Likewise, neither *smax1 smxl2* nor *smxl6,7,8* displayed altered flg22-induced resistance. Together, these results suggest that both KL and SL signalling contribute to MAMP-triggered ROS production, while the KAI2-MAX2 module plays a more prominent role in PTI by further affecting flg22-induced resistance, potentially through mechanisms independent of SMAX1, SMXL2 and SMXL6,7,8.

### Karrikin and strigolactone signalling modulate the strength of the transcriptional response to PTI activation

To investigate transcriptional changes connected to the altered PTI-associated ROS bursts observed in KL- and SL-signalling mutants, we performed transcriptome profiling of adult leaves from wild type and mutant plants infiltrated with water (mock), flg22 or chitin. To capture temporal dynamics of immune activation, samples were harvested at 30- and 180-minutes post infiltration (mpi).

Prior to analysis of differentially expressed genes, samples were inspected through hierarchical clustering of sample distance and principal component analysis. Three samples were deemed outliers based on hierarchical clustering of sample distance and were removed before downstream analysis (Fig. S4A).

Principal component analysis revealed strong clustering primarily according to the timing of treatment, indicating that temporal progression of the immune response, rather than genotype, accounted for the largest proportion of transcriptional variation across samples (Fig. S4B). When samples collected at the two time points were analysed separately, elicitor-treated samples separated clearly from mock-treated controls at 30 mpi, whereas flg22- and chitin-treated samples largely overlapped (Fig. S4C). By contrast, transcriptional responses to flg22 and chitin diverged more strongly at 180 mpi, indicating increasing elicitor-specific signalling over time (Fig. S4D). Consistent with this observation, early responses at 30 mpi included a shared set of 258 differentially expressed genes (DEGs) common to both flg22- and chitin-treated leaves across all genotypes (Fig. 3A). At 180 minutes, only flg22-treated leaves exhibited a shared group of 196 DEGs across genotypes (Fig. 3B). Notably, an additional set of 668 DEGs was shared among all genotypes after flg22 treatment, except *d14-1*, raising the question of a reduced transcriptional response to flg22 in this mutant (Fig. 3B).

**Figure 3:**
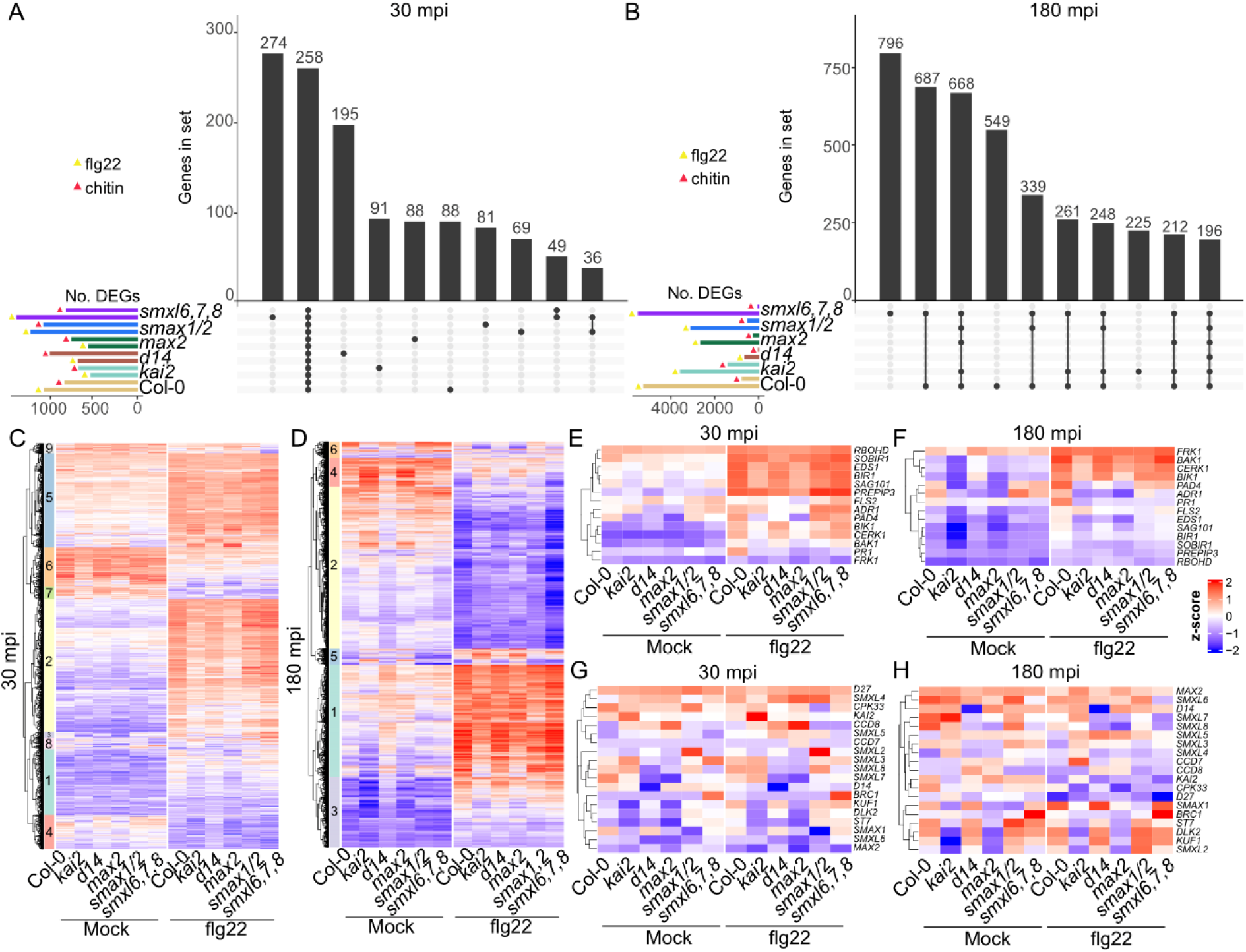
The amplitude of the transcriptional response to flg22 and chitin is modulated in adult leaves of KL and SL signalling mutants. A-B) Upset plots comparing the sets of differentially ex-pressed genes (DEGs; |Log2FC| > 1 & padj < 0.05) in flg22 or chitin vs mock treated leaves across genotypes **A)** 30 min or **B)** 180 min after infiltration. Vertical black bars and numbers correspond to the number of genes that are present in sets indicated at the x-axis, and connected dots below the bar indicate in which genotype and treatment combination these genes are differentially expressed. **C-D)** k-means clustered heatmaps of all DEGs in response to flg22 treatment across genotypes at **C)** 30 mpi (1831 DEGs) or **D)** 180 mpi (7070 DEGs), depicted as relative expression (z-score). These clusters were used for Gene Ontology enrichment in supplementary Figure S8. Hierarchically clustered heatmaps showing expression levels (z-score) of genes associated with (**E-F)** pattern- and effector-triggered immunity, and (**G-H**) karrikin or strigolactone signalling at 30 mpi and 180 mpi with and without flg22 treatment.

Across both timepoints, Col-0 plants exhibited a larger number of DEGs in response to flg22 compared to chitin treatment, consistent with previous observations in MAMP-treated Arabidopsis seedlings (75) (Fig. 3A, B). This difference became more pronounced at 180 mpi, when flg22 continued to induce strong transcriptional reprogramming while chitin responses had largely subsided (Fig. 3B). Accordingly, all genotypes displayed stronger late transcriptional responses to flg22 than to chitin (Fig. 3B). At 30 mpi, however, *kai2-2*, *d14-1*, and *max2-1* mutants exhibited fewer flg22-responsive DEGs compared to the other genotypes (Fig. 3A). This reduction paralleled the impaired MAMP-triggered ROS burst observed in these mutants (Fig. 2A-D). Consistently, leaves of *smxl6,7,8*, which had displayed enhanced ROS production, showed the largest number of flg22-responsive DEGs at both timepoints (Fig. 2E; 3A, B), supporting a link between ROS amplitude and transcriptional responsiveness.

Overall, at 30 mpi, the different genotypes show a stronger variation in number of DEGs in response to flg22 treatment, whereas there was less of a genotype effect on the number of DEGs in response to chitin treatment (Fig. 3A). Additionally, we compared sets of DEGs across genotypes compared to Col-0 under mock conditions at either 30 or 180 mpi (Fig. S5A, B). At both timepoints a number of DEGs were shared between *kai2* and *max2* (174 DEGs at 30 mpi, and 515 DEGs at 180 mpi), indicating that these DEGs are most likely related to perturbed KAR/KL signalling (Fig. S5A, B). Surprisingly, only few DEGs were observed in both *smax1 smxl2* and *smxl6,7,8* compared to Col-0 under mock conditions, at either timepoint (Fig. S5A, B).

To uncover any contrasting transcriptional responses to flg22 or chitin across genotypes, we compiled all respective DEGs in response to flg22 or chitin treatment across genotypes per harvest time point and visualized the relative expression value (z-score) and log2 fold change (log_2_FC) in heatmaps upon k-means clustering and Gene Ontology enrichment (Fig. 3C-D; S6-S8). GO term enrichment analysis revealed that clusters showing strong gene expression changes in response to flg22-treatment were enriched for immune-response processes associated with defence against bacteria (Fig. S8). In contrast, clusters of strongly downregulated genes were enriched for photosynthesis- and thylakoid-related genes, potentially indicating a metabolic shift from CO_2_ fixation and growth to defence during immune activation (Fig. S8).

Overall, little to no genes responded in a contrasting manner to flg22 or chitin across genotypes (Fig. 3C, D; S6, S7). However, expression amplitudes in flg22-responsive gene clusters were highest in Col-0 and *smxl6,7,8* plants, and consistently weaker in *kai2-2* and *max2-1* mutants (Fig. 3C, D; Fig. S6). This correlates with the variations in PTI-triggered ROS responses we found across the mutants, and suggests that KAI2-MAX2- and SMXL6,7,8-mediated signalling modulates the strength rather than the identity of PTI (transcriptional) outputs. Although distinct from *kai2-2* and *max2-1*, *d14-1* leaves also exhibited weaker expression amplitudes of flg22-induced genes at 180 mpi compared to Col-0, (Fig. S6; S9B). Although *kai2-2, d14-1* and *max2-1* show similarly impaired MAMP-triggered ROS production, their transcriptional responses to flg22 treatment are differentially affected (Fig. 3C-H; 2; S9).

We compared the relative expression of canonical immune regulators and found that baseline expression levels of several defence-associated genes, including *ENHANCED DISEASE SUSCEPTIBILITY 1* (*EDS1*), *PHYTOALEXIN DEFICIENT 4* (*PAD4*), *SENESCENCE-ASSOCIATED CARBOXYLESTERASE 101* (*SAG101*), and *ACTIVATED DISEASE RESISTANCE 1* (*ADR1*), differed among genotypes at 180 mpi, but elicitor-induced fold changes were largely comparable (Fig. 3E, F). This suggests that the total abundance of defence-related transcripts are affected in our mutants, but the relative changes upon elicitation are similar to Col-0. Only *PATHOGENESIS RELATED 1* (*PR1*) displayed reduced flg22-induced expression in *kai2-2* and *max2-2* compared to the other genotypes, both in terms of relative expression and fold changes compared to mock treated samples at 30 and 180 mpi (Fig. 3E, F; S9A, B). Together, these results indicate that KL and SL signalling mutants primarily differ in basal transcriptional states translating into lower relative abundance upon treatment, rather than in their general capacity to activate core PTI transcriptional programs (Fig. 3C-F; S9A, B).

### *KUF1* is induced by MAMPs independently of KAI2 and MAX2

To understand whether flg22 or chitin activates KL or SL signalling, we analysed the expression and fold-change of KL and SL signalling related genes upon mock or flg22 treatment (Fig. 3G, H; S9D, E). The KL-responsive genes *DWARF14-LIKE2 (DLK2*) and *KARRIKIN UPREGULATED F-BOX1* (*KUF1*), which are transcriptionally induced upon activation of KAI2-mediated signalling, showed reduced basal expression in *kai2-2* and *max2-1* mutants and increased basal expression in *smax1 smxl2* compared to the wild type, consistent with previous reports (Fig 3G, H) (6,15,18). Expression of *KUF1* was upregulated by flg22 treatment both at 30 and 180 mpi in all genotypes, with lower relative expression in *kai2-2* and *max2-1* (Fig. 3G, H; S9D, E). *KUF1* expression was only significantly induced at 30 mpi with chitin (Fig. S9D, E). Unexpectedly, fold change induction of *KUF1* upon flg22 treatment in both *kai2-2* and *max2-1* was comparable to wild type (especially at 180 mpi), indicating that immune activation can either bypass canonical KL-signalling that normally restricts *KUF1* induction during treatments with KARs or the synthetic SL GR24 (Fig. S9C, D) (8,18), or that *KUF1* may be induced by independent pathways. In contrast, *DLK2* and the SL marker gene *BRANCHED1* (*BRC1*) were not induced in Col-0 by flg22, but at basal levels *smxl6,7,8* plants exhibited the highest relative expression of *BRC1*.

Overall, transcriptome profiling showed that both the *kai2-2* and *max2-1* mutants cluster separately from other genotypes following elicitor treatment, despite exhibiting qualitatively similar gene expression changes (Fig. S4C;3C-H). Combined with ROS phenotypes observed in *kai2-2*, *max2-1*, and *d14-1* mutants, these results suggest that reduced PTI outputs likely arise from multiple regulatory mechanisms rather than a single transcriptional defect (e. g. due to reduced ROS production).

### KL and SL signalling affect genes involved in phytohormone biosynthesis and catabolism

KL- and SL-signalling are known to interact with several other hormonal pathways, including ABA (9), auxin and ethylene (12,13,44), as well as SA signalling (67,76,77). To assess whether potential interactions are reflected in our transcriptomic dataset, we examined transcript abundance of genes associated with the biosynthesis and signalling of JA, SA, ABA and ethylene (Fig. S10-S13).

In *kai2-2* and *max2-1* several SA-signalling, -biosynthesis and -marker genes displayed reduced basal expression and also lower expression after flg22 and chitin treatment at 30 and 180 mpi (Fig. S10A, B). Fold change values in response to flg22 or chitin treatment, however, were comparable across genotypes (Fig. S10C-E). MAMP-induced expression of JA-associated genes was largely comparable among genotypes (Fig. S11A-D). Interestingly though, basal expression of JA-associated genes, including the biosynthesis enzyme encoding genes *ALLENE OXIDASE SYNTHASE* (*AOS), ALLENE OXIDASE CYCLASE 1 (AOC1)*, were enhanced in *kai2-2, d14-1* and *max2-1* at 180 mpi, but reduced in both *smax1 smxl2* and *smxl6,7,8*, compared to Col-0 (Fig. S11E). Expression of ABA and ethylene biosynthesis and response genes was largely comparable across all genotypes, both in terms of relative expression, basal expression and fold change values (Fig. S12, S13). This suggests a predominant role of the SL and KL pathway components in the modulation of both JA and SA biosynthesis and signalling.

### Modulation of PTI by KAI2- and D14- signalling is conserved across dicotyledons

*KAI2* genes are present throughout land plants and Charophyte algae, whereas the SL receptor gene *D14* emerged in basal embryophytes through a gene duplication event of ancient *KAI2* (35,36). Consistent with this evolutionary conservation, developmental and symbiosis-related functions of KAI2- and D14-signalling have been reported across diverse plant families, including *Poaceae* (40,45,47,48,78) *Fabaceae* (50,52), *Solanaceae* (33,79) and *Brassicaceae* (11,28). This raises the question, whether the regulatory role of these receptors in immune responses is likewise conserved.

To address this, we analysed KL- and SL- mutants in selected plant species for altered ROS production upon MAMP treatment. The legume *Lotus japonicus*, a model species for studying symbiotic interactions with arbuscular mycorrhizal fungi and nitrogen-fixing rhizobia, has predominantly been investigated in the context of root-associated ROS responses (80). As the pathogens examined in this study primarily infect aerial tissues, we assessed ROS production in leaves by challenging leaf discs of mature plants with flg22. Similar to our observations in Arabidopsis, *kai2a-1 kai2b-1*, *d14-1* and *max2-3* mutants showed reduced flg22-triggered ROS production compared to the wild type (Fig. 4A). However, flg22-induced ROS production was not affected in the *smax1-3* mutant (Fig. 4A). We also examined the *Nicotiana benthamiana KAI2* quadruple mutant *kai2a,b,c,d* for altered MAMP-triggered ROS production. Consistent with our results from Arabidopsis and *L. japonicus*, the *N. benthamiana* quadruple mutant showed strongly reduced flg22- and chitin-triggered ROS production (Fig. 4B, C). Collectively, these findings indicate that KAI2- and D14-mediated regulation of PTI signalling represents a conserved im-mune-modulatory mechanism across diverse dicotyledonous lineages.

**Figure 4:**
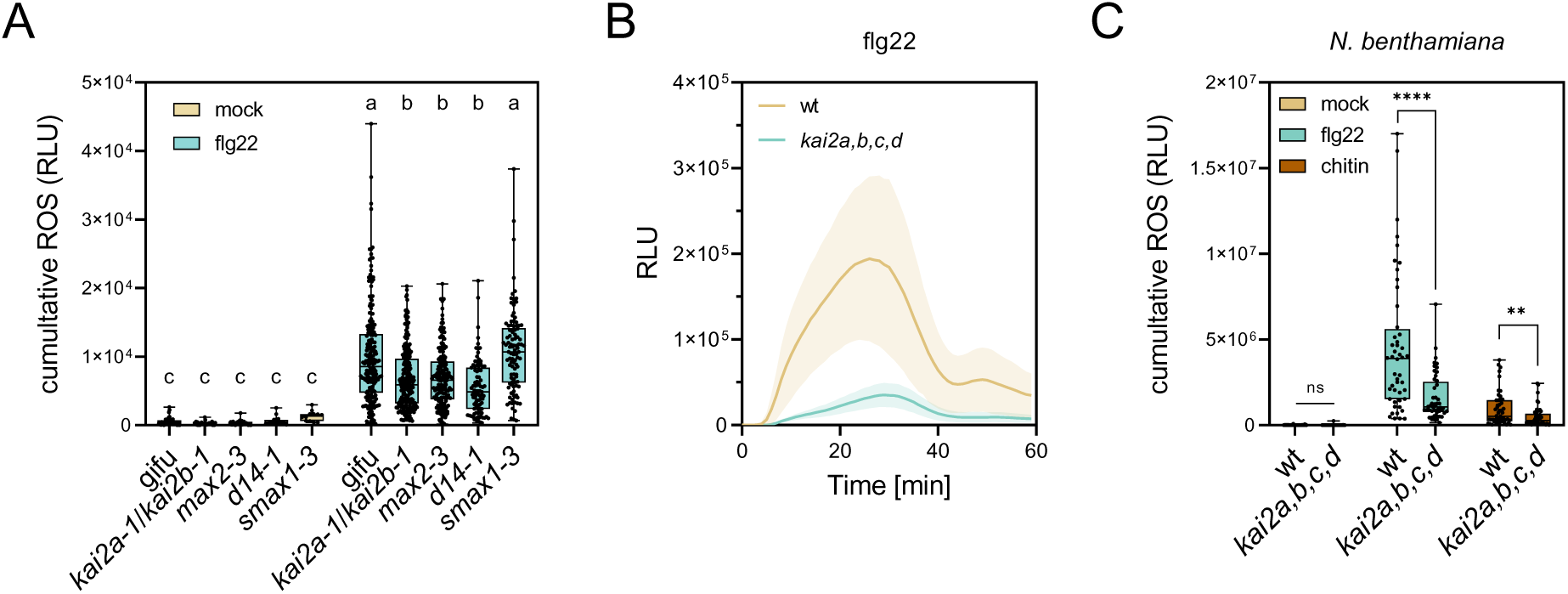
Regulation of flg22-triggered ROS by KAI2 and D14 extends to other dicotyledonous species. ROS productioni in leaf discs of the indicated genotypes as relative light units (RLU). **A**) Total RLU after elicitation with flg22 (1 µM) in *L. japonicus*; mean ± SD, n ≥ 35, with data pooled from six independent replicates (one-way ANOVA, Tukey’s post-hoc test; a-b/c: p < 0.0001, b-c: p ≤ 0.01). **B**) ROS kinetics of *N. benthamiana* leaf discs after elicitation with flg22 (100 nM); mean ± SD, n = 12. **C)** Total RLU after elicitation with flg22 (100 nM) or chitin (20 ng/µl) in *N. benthamiana*; mean ± SD, n = 48, data pooled from four independent replicates (two-tailed Student’s t-test; **: p < 0.005, ****: p < 0.0001).

## Discussion

Multiple phytohormones play central roles during plant adaptation to biotic stress (2). Our work identifies a new role for the KL and SL signalling modules within this regulatory network. Disruption of both KL and SL receptor complexes resulted in a strong reduction of MAMP-triggered ROS production, while effects on MAMP-triggered transcriptional responses and resistance against pathogens were specific to KL receptor components. Surprisingly, *smxl6,7,8* displayed enhanced ROS production and stronger transcriptional responses to flg22 and chitin. By contrast, *smax1 smxl2* showed ROS and transcriptional responses largely comparable to wild type. The opposing ROS and transcriptional phenotypes observed between *smxl6,7,8* and receptor complex mutants support a potentially unconventional mechanistic interaction between the KAI2-MAX2 module and SMXL6,7,8 transcriptional repressors during PTI. KAI2-MAX2 primarily interacts with and targets SMAX1 for MAX2-dependent proteasomal degradation, while SMXL6,7,8 are primarily targeted by the D14-MAX2-signalling module (20,37). Although both pathways share MAX2, non-canonical SMXL targeting has previously primarily been reported for D14 (25). However, recent research demonstrated that the SMAX1 and the SMXL68 clade are not exclusively targeted by the KAR/KL or SL pathway, respectively. Both SMAX1 and SMXL2 can be ubiquitylated and de-graded after SL recognition in a D14-dependent manner, demonstrating cross-regulation at the level of SMAX/SMXLs between the KAR and SL pathway (24,25). Moreover, the previously considered SL-specific SMXL6,7,8 interact with DDB1-BINDING WD-REPEAT DOMAIN HYPERSENSITIVE TO ABA DEFICIENT1 (DWA1), conferring substrate specificity to a CULLIN4 (CUL4)-type E3 ubiquitin ligase, which is involved in translational de-repression of ABA- and drought responsive genes (81). These findings support the suggested existence of a MAX2-independent mechanism for SMXL turnover (23). Furthermore, genetic studies imply SMXL6,7,8 as potential growth-condition-dependent targets of KAI2 mediated turnover, as *smxl6,7,8* mutants showed similar root skewing phenotypes to *smax1 smxl2* mutants and opposite phenotypes to *kai2* and *max2* mutants (11,12). Our data further support a potential mechanistic connection between SMXL6,7,8 and KAI2-MAX2 signalling during PTI, but the mechanistic basis remains unknown.

Our transcriptome data suggest that KL signalling primarily influences the amplitude of PTI rather than its qualitative identity. SA signalling represents a central determinant of resistance against (hemi-)biotrophic pathogens (82) and has previously been linked to immune phenotypes in KL receptor mutants (76). Earlier studies reported reduced basal expression of SA-associated genes in KL receptor mutants together with initially delayed, yet ultimately stronger, SA induction after 6 hours of *Pto* infection (76). Consistent with this observation, our transcriptomic data reveals reduced basal expression of SA and JA biosynthesis and marker genes in KL signalling mutants (Fig. S10, S11). Across wild type and the mutants, fold change in expression of SA- and JA- related genes were comparable between mock and MAMP treatments (Fig. S10, S11). This raises the question whether the difference across basal expression explains the reduced PTI phenotypes of KL signalling mutants.

Our data show that flg22-induced resistance was compromised in both *kai2-2* and *max2-1* (Fig. 2G). This suggests a functional contribution of KL signalling to PTI-induced resistance. Interestingly, MAPK activation remained unaffected across the analysed genotypes, indicating that disruption of KL- or SL-signalling selectively affects distinct PTI-associated outputs (Fig. S3). This is consistent with previous findings, showing that MAPK activation and ROS production are mechanistically uncoupled PTI outputs. For example, RBOHD, required for MAMP-triggered ROS production, does not contribute to flg22-induced MAPK activation (83). Interestingly, ROS is important for long distance communication during stress responses (84–86). This raises the question whether the role of MAX2-function in systemic acquired resistance is associated with defects in stimulus-triggered ROS production (67,77). Previous studies also reported a link between increased bacterial pathogen susceptibility of *max2* to altered stomatal regulation and ROS sensitivity (66), further corroborating our findings of ROS-related PTI signalling defects of KL signalling mutants.

Despite the general involvement of KL and SL signalling components in PTI, their influence on pathogen resistance was diverse (Fig. 1). Mutants defective in KL signalling (*kai2* and *max2*) displayed enhanced resistance to *Ecr* relative to Col-0 (Fig. 1A, B). Surprisingly, *smxl6,7,8* mutants showed a comparable *Ecr* resistance phenotype, raising the question whether their role for *Ecr* accommodation is independent of their canonical mutual cross-regulation during SL/KL signalling. These findings indicate that KL-related signalling components can function as susceptibility factors for certain obligate biotrophic pathogens. It remains unclear how reduced PTI responses in *kai2* and *max2* are correlated with enhanced resistance to powdery mildew. It is possible that powdery mildew hijacks a yet elusive endogenous KL-dependent pathway for its own benefit. KL signalling is important for accommodation of AM fungi, raising the question whether KL signalling may be hijacked by powdery mildew. Interestingly, mutants in the receptor kinase FERONIA also display reduced PTI responses, but increased resistance to powdery mildew fungi (87–89). However, whether both pathways converge remains elusive.

By contrast, constitutive de-repression of KL and SL signalling in *smax1 smxl2* and *smxl6,7,8* mutants resulted in increased susceptibility to necrotrophic fungal pathogens (Fig. 1D; S1B). Reduced basal JA biosynthesis in these mutants may compromise defence against necrotrophs, providing a potential explanation for their enhanced susceptibility. Interestingly, this contrasts with observations in tomato, where SL-deficient plants display increased susceptibility to *A. alternata* and *B. cinerea*, raising the question of species-specific differences in the integration of SL signalling with immune networks (90). Conversely, *smax1 smxl2* and *smxl6,7,8* mutants resulted in increased resistance to biotrophic *Hpa*, suggesting that SMXL-mediated regulatory mechanisms contribute to host susceptibility in this interaction (Fig. 1C; S1A).

Importantly, the involvement of KL and SL signalling in the regulation of PTI responses appears evolutionary conserved in dicotyledonous species, including *L. japonicus* and *N. benthamiana*. KAI2 is evolutionarily ancient and appeared in green algae, while emergence of the SL receptor D14 occurred via a KAI2 gene duplication event in angiosperms (35,36). This suggests that regulation of MAMP-triggered ROS responses may have occurred at latest in the last common ancestor of dicotyledons. It will be interesting to test whether this regulatory function can be found in monocotyledons and further traced back in the green lineage. Throughout land plant evolution the KAR/KL signalling pathway has changed to regulate distinct biological processes. For example, KL signalling contributes to the regulation of SL biosynthesis in AM-competent angiosperms but not in Arabidopsis or liverworts, that lost KL signalling target genes involved in AM fungal accommodation (44,46,91,92). Notably, Arabidopsis *kai2-2* mutants shared more transcriptional similarities with *max2-1* than with *d14-1*, suggesting that KAI2-dependent signalling represents a major branch of MAX2-mediated signalling in Arabidopsis leaves. This observation is consistent with the earlier emergence of KAI2 relative to D14 and a longer co-evolutionary history between KAI2 and MAX2 (16,35,36)

Collectively, our findings identify KL and SL signalling as conserved modulators of plant immunity that do not influence the identity but the amplitude of early PTI responses and alter pathogen resistance. The integration of these signalling pathways that are also involved in regulation of plant development, with immune and hormonal networks highlights the complexity of regulatory mechanisms balancing growth, symbiosis, and defence in plants. The mechanistic basis of KL and SL-dependent modulation of PTI and pathogen infection outcomes remains to be determined in future studies.

## Materials and Methods

### Plant material and growth conditions

Arabidopsis seeds were stratified on soil for 2 - 3 days in the dark at 4 °C prior to transfer to growth conditions. For soil-based assays (CL ED73, Einheitserde(R)), two plants per pot were cultivated in controlled environmental conditions (20/18 °C, 60% relative humidity, 8 h photoperiod) for 5 - 6 weeks. *Nicotiana benthamiana* plants grown in controlled environmental conditions with one plant per pot (21 °C, 60% relative humidity, 16 h photoperiod) without stratification. *Lotus japonicus* seeds were sterilised with 1% NaClO and 0.1% SDS and then washed five times with MilliQ water. Imbibed seeds germinated on 0.8% plant agar plates. 10-day-old seedlings were initially planted in soil (CL ED73, Einheitserde(R)), and consequently moved to a mixture of potting soil and quartz sand (1/2 sand/soil; potting Einheitserde).

### Detection of ROS accumulation

For Arabidopsis and *N. benthamiana*, leaf discs (⌀ 4 mm) from five- to six-week-old plants were incubated floating on water overnight in white 96-well plates in the same temperature and light conditions in which the plants were grown. The water was replaced by a solution containing 2 µg/ml horseradish peroxidase and 5 µM L-012. Emitted luminescence was recorded as relative light units (RLU) in 1 min intervals after 10 min background reading followed by elicitor treatment for 1 h. For *L. japonicus*, leaf discs (⌀ 3 mm) were taken from four- to five-week-old plants. The same procedure was followed for Arabidopsis and *N. benthamiana*. The substrate solution for ROS detection in *L. japonicus* leaf discs contained 40 nM L-012 and 8 ng/ml HRP. Luminescence values were normalised to average background luminescence per well.

### Analysis of flg22-induced resistance

Leaves of five- to six-week-old Arabidopsis plants were syringe infiltrated with either mock or flg22 24 h prior to infection. *Pseudomonas syringae pv. tomato* DC3000 was grown on King’s B agar plates containing 50 μg/ml rifampicin at 28°C. After two-to-three days, bacteria were resuspended in ddH_2_O containing 0.04% Silwet L77. The bacterial suspension was adjusted to an OD_600_= 0.0002 (=1 x 10^5^ cfu/ml) before syringe infiltrating pre-treated leaves. Bacterial counts were determined two days after infection by re-isolation and plating of colonies with subsequent colony counting. Bacterial growth was calculated as cfu/cm^2^ of leaf area.

### MAPK activation and Western blot analysis

Five- to six-week-old Arabidopsis plants were used for MAPK-assay. One leaf each from four plants was syringe infiltrated per timepoint with either mock or 1 µM flg22. Samples were frozen in liquid nitrogen and homogenized using a tissue lyser. Proteins were extracted using a 50 mM Tris-HCl (pH 7.5) buffer containing 50 mM NaCl, 10% glycerol, 5 mM DTT, 1% protease inhibitor cocktail, 1 mM phenylmethylsulfonyl fluoride (PMSF), 1% IGEPAL, 10 mM EGTA, 3 mM NaF, 3 mM Na3VO4, 4 mM Na2MoO4, 30 mM ß-Glycerophosphate and 30 mM p-nitrophenylphosphate before analysis by SDS-PAGE and Western blot. Phosphorylated MAPKs were detected using 1:2000 α-p44/42 primary antibodies (Cell Signaling Technology, Cat. 9101S) and 1:10.000 secondary mouse α-rabbit IgG-HRP (Santa Cruz, sc-2357) antibodies. Band intensities of MPKs were quantified using ImageJ and normalised to the Rubisco band (CBB stain) for each lane. Normalised values were expressed relative to the calibrator sample which was set to 1 and is indicated in the respective figure legend.

### Detection of protein accumulation by Western Blot

Leaves of five- to six-week-old Arabidopsis plants were harvested and total protein extracted with a 50 mM Tris-HCl (pH 7.5) buffer containing 50 mM NaCl, 10% glycerol, 2 mM EDTA, 2 mM DTT, 1% protease inhibitor cocktail, 1 mM phenylmethylsulfonyl fluoride (PMSF) and 1% IGEPAL. Analysis was performed by SDS-PAGE and Western blot, detecting protein accumulation with specific α-RBOHD (1: 500, Agrisera, AS15 2962), α-FLS2 (1:2000, Agrisera, AS12 1857) and α-CERK1 (1:2000, Agrisera, AS16 4037) and secondary mouse α-rabbit IgG-HRP antibodies (Santa Cruz, sc-2357). Band intensities were quantified using ImageJ and normalised to the Rubisco band (CBB stain) for each lane. Normalised values were expressed relative to the calibrator sample which was set to 1 and is indicated in the respective figure legend.

### RNA-sequencing and data analysis

Leaves of five- to six-week-old Arabidopsis plants were syringe infiltrated with either mock, 1 µM flg22 or 75 µg/ml chitin for 30 min or 180 min. One leaf each from four plants per genotype was harvested, ground and total RNA isolated using the Spectrum Plant Total RNA Kit (Sigma-Aldrich, USA). RNA was sent to Novogene for paired-end sequencing, with sequencing of up to 20 million reads per sample. Reads were mapped to Araport 11 using Salmon as previously described in (93). Mapped reads were imported into an R studio pipeline using *Tximport.* We used DESeq2 (94) to identify differentially expressed genes (|Log_2_FC| ≥ 1; padj < 0.05) after treatment as well as create to a relative expression matrix using a variance stabilizing transformation (VST). Heatmaps based on relative expression values or Log_2_FC were plotted using the *ComplexHeatmap* package, and the number of clusters for K-means clustering was decided after inspecting silhouette and elbow scores. Subsequent GO term enrichment analysis was performed using the *clusterProfiler* package.

### Powdery mildew propagation, plant infection and conidiophore quantification

Powdery mildew fungi of the species *Erysiphe cruciferarum* (*Ecr*) were propagated weekly by transferring spores to four- to five-week-old uninfected Col-0 and *pad4* plants. Powdery mildew spores were used for further inoculation 2 - 3 weeks after propagation. Powdery mildew propagation and infection experiments were performed in an environmentally controlled growth cabinet (22 °C, 70% relative humidity, 8 h photoperiod). Conidiophore production was analysed 5 days after infection. To this end, four leaves per genotype were harvested from individual infected plants. The leaves were destained using EtOH:Acetic Acid (6:1) and subsequently stained using ink (ink:25% acetic acid; 9:1). Ink-stained fungal structures were visualized using the Zeiss AXIO Lab microscope with a 20x magnification for conidiophore counting.

### Infection with *Hyaloperonospora arabidopsidis* (*Hpa*) and spore quantification

For *Hpa* infection assays, seeds were stratified on soil for 2 - 3 days in the dark at 4 °C and grown in environmentally controlled conditions (22 °C, 60% relative humidity, 16 h photoperiod). 12-day-old seedlings were spray-inoculated with a spore suspension of *Hpa* isolate Noco2 (5 × 10^4^ spores/ml) (95). After inoculation, plants were kept in trays covered with sealed lids to maintain high humidity (18 °C, near-saturation humidity). At 6 days post inoculation, infected aerial parts were harvested in water and vortexed to release spores. Spores were counted using a hemocytometer and normalised to leaf fresh weight. For imaging, cotyledons or leaves were harvested and stained in 0.01% (w/v) trypan blue in lactophenol for 3 min at 95 °C followed by incubation for 5 h at room temperature. Samples were subsequently cleared overnight in chloral hydrate (2.5 g/ml) and mounted in glycerol. Pictures were obtained using the Zeiss Apotome II system.

### Infection with *Alternaria brassicicola* and *Botrytis cinerea* and scoring of disease symptoms

*A. brassicicola* and *B. cinerea* spores were diluted with water to a final density of 5 x 10^5^ spores/ml. Plants were inoculated by placing two 5 µl droplets of spore suspension onto the leaf surface. Inoculated plants were kept at 100% relative humidity at 22 °C at an 8 h light/16 h dark cycle. Growth of *A. brassicicola* was scored after 13 days by classifying symptom severity ranging from 1 (no symptoms), 2 (light necrotic lesions), 3 (severe necrotic lesions), 4 (spreading of lesions beyond infection site), 5 (whole leaf affected) to 6 (sporulation of the fungus). A disease index (DI) was calculated using the following formula: DI = S_i_ x n_i_; S_i_ is the symptom category, n_i_ is the percentage of leaves in i. Pictures of detached leaves were taken after 13 days. *Botrytis cinerea* infection was scored by measuring the lesion size at the inoculation site with ImageJ on images taken after 4 days.

### Viral infection assays

Viral infection assays were performed as previously described (96). In short, Arabidopsis seeds were germinated on half-strength Murashige and Skoog (MS) medium and transplanted to soil after one week.

Plants were grown in a controlled growth chamber under long-day conditions (16 h light/8 h dark) at 24°C. *Agrobacterium tumefaciens* strain GV3101 carrying pBin1.2_BCTV (97) or pBIN-plus vector with a C-terminal GFP tag (98) was grown on Petri dishes with LB agar supplemented with appropriate antibiotics. Bacterial cells were scraped from plates and inoculated into the centre of the rosette of 2- to 3-week-old plants by mechanical wounding using an entomology pin, with 20 punctures per plant. Samples were collected at 21 days post-inoculation (dpi). Total DNA was extracted using a CTAB-based method (99) from infected flower tissues or systemic leaves, and viral DNA accumulation was quantified by qPCR using primers targeting BCTV Rep gene (Fw: ATGCAAGAATGGGCTGATGC, Rv: TCTGCCAC-TCCTTTTGTGCT), normalised to *At*Actin2 (Fw: CAGTGTCTGGATCGGTGGTT, Rv: TGAACGAT-TCCTGGACCTGC). The reactions were performed as follows: 2 min at 95°C, 40 cycles consisting of 15 s at 95°C, 1 min at 60°C. The experiment was independently repeated four times with each treatment including at least nine plants per experiment. Each sample was analysed in three technical replicates for qPCR. Relative viral DNA levels were calculated using the 2^−ΔΔCt^ method.

### Elicitors

The flg22 peptide (QRLSTGSRINSAKDDAAGLQIA) was synthesized by Biomatik (Canada), elf18 (SKEKFERTKPHVNVGTIG) by Pepmic (China) and *Fo*nlp20 (AIMYAWYWPKDQPADGNLVSGHR), derived from *Fusarium oxysporum*, was kindly provided by Dr. Justin Lee (IPB Halle). Shrimp shell-derived chitin (Sigma-Aldrich, Germany) was ground and extracted with ddH_2_O. Concentrations refer to µg chitin powder dissolved per ml of water before extraction.

### Statistical analysis

GraphPad Prism (Version 8.0.1) was used to perform statistical analysis (except for RNA-seq data analysis). Detailed descriptions of the sample size, p-values and statistical methods are indicated in the respective figure legends.

## Acknowledgements

We thank Franziska Brückner (Max Planck Institute of Molecular Plant Physiology) for extracting RNA for transcriptomics, and Iván F. Acosta (Max Planck Institute of Molecular Plant Physiology) for help with transcriptomics analysis.

## Author contributions

Conceptualization: MS, CG; Investigation: SDS, KB, AS, DM, AM, HW. Funding acquisition: MS, CG, RL-D, MKR-L, BK. Project administration: MS, CG; Supervision: MS, CG, RL-D, MKR-L, BK, SDS. Writ-ing - original draft: SDS, KB. Writing - review & editing: SDS, KB, CG, MS with input from all authors

## Supplemental Figures

**Figure S1:**
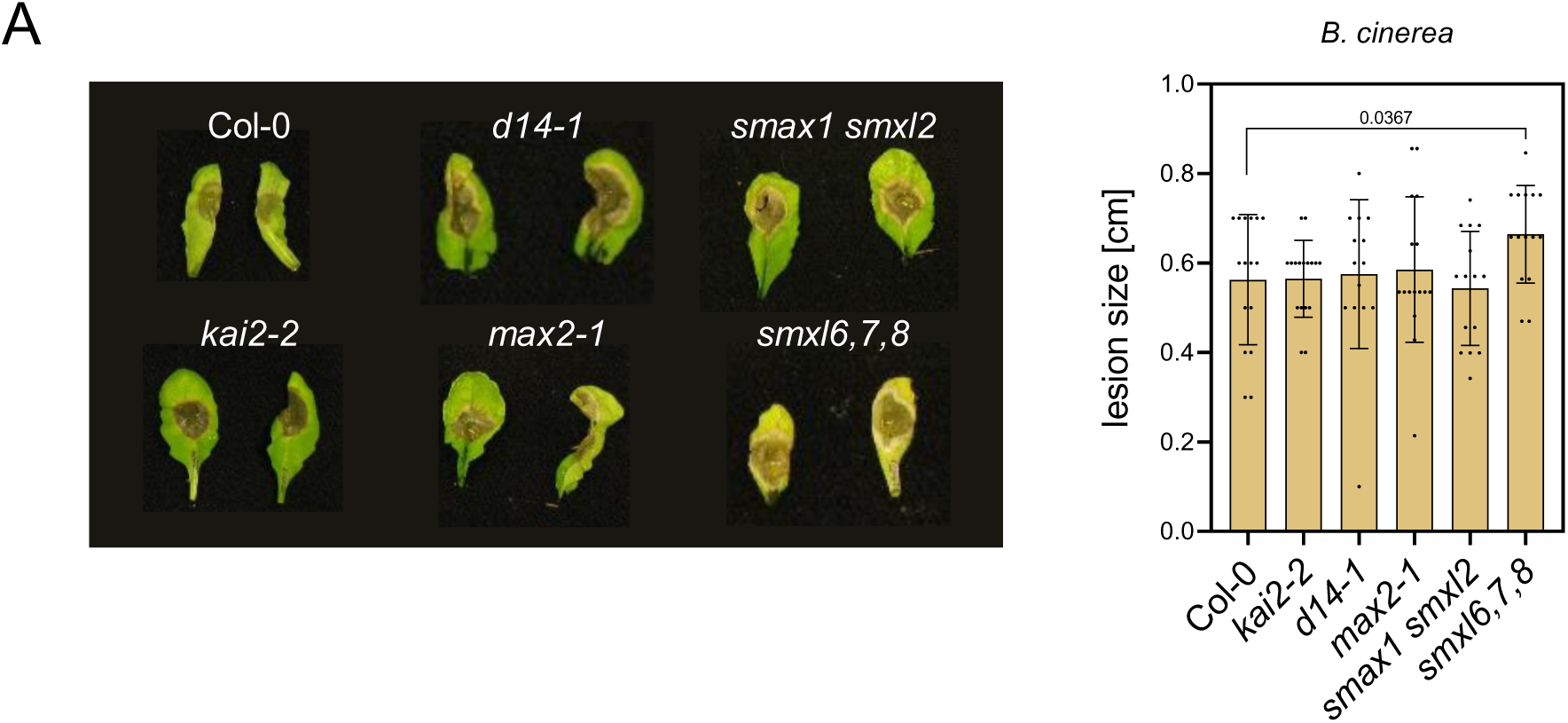
Increased susceptibility of *smxl6,7,8* mutants to *B. cinerea*. **A)** Representative images of detached infected leaves of the indicated genotypes. Lesion size of *B. cinerea* infected leaves 4 days post-infection. Mean ± SD, n = 14 - 17, pooled from three independent experiments (Student’s t-test; p-value is indicated in the figure).

**Figure S2:**
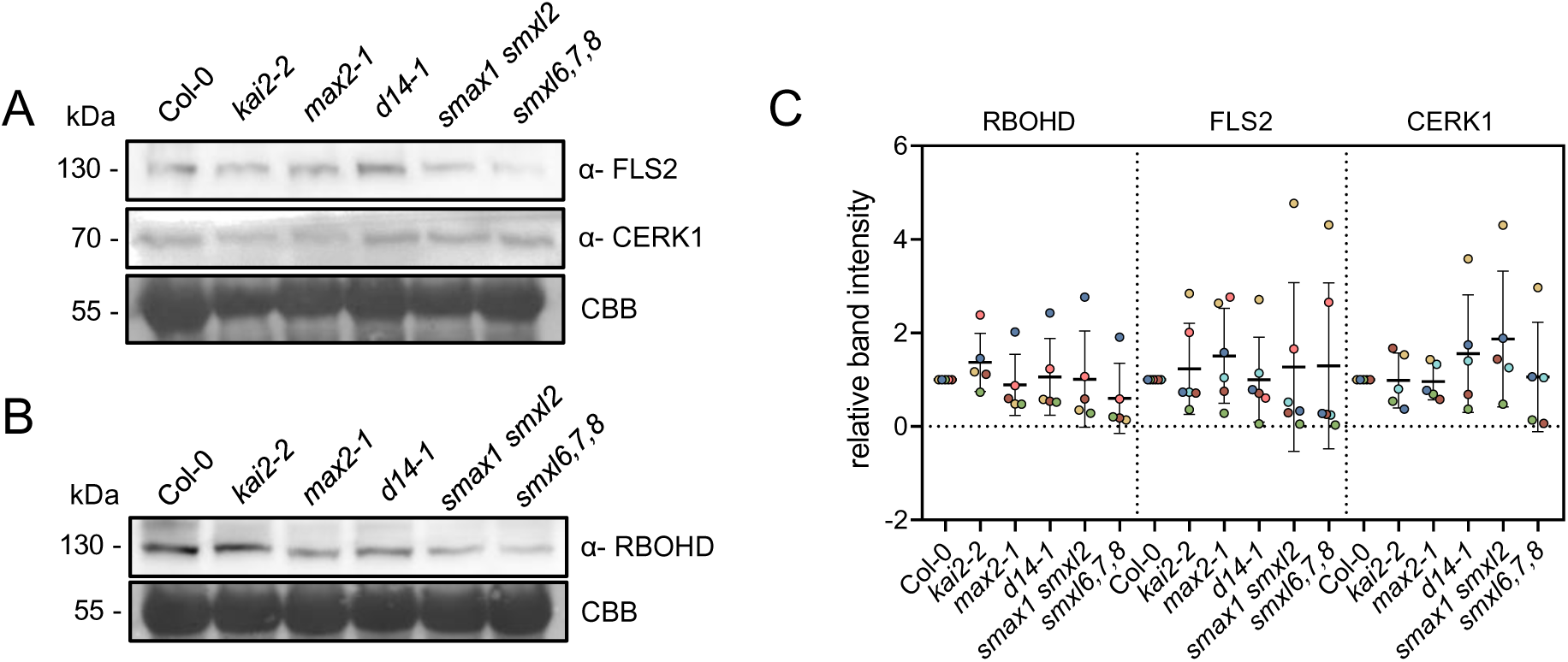
Protein accumulation of FLS2, CERK1 and RBOHD in KL and SL signalling mutants. A-B) Total protein of mature plants assayed by Western Blot using specific antibodies raised against **A)** FLS2 and CERK1 or **B)** RBOHD. CBB = Coomassie Brilliant Blue. **C)** Relative band intensities detected in **A)** and **B)**. Band intensities of six experiments were measured using ImageJ software and normalized against CBB and relative to Col-0. Data points indicate individual experiments with points from the same experiment being marked with the same color.

**Figure S3:**
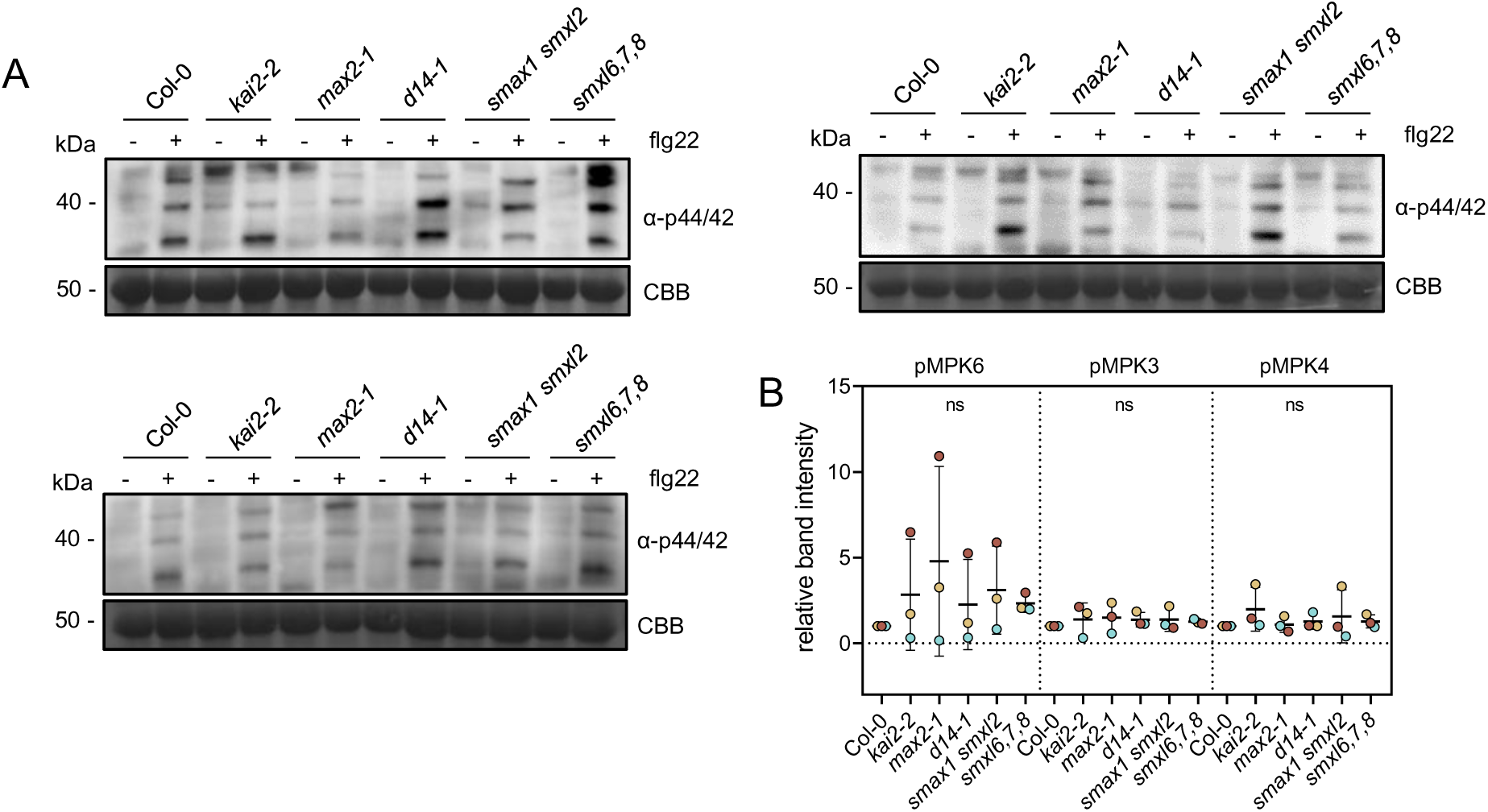
flg22-triggered MAPK activation is not altered in KL and SL signalling mutants. MAPK activation in indicated genotypes 20 min after syringe-infiltration of 1 µM flg22 into mature rosette leaves. Western blots were probed with α-p44/42. CBB = Coomassie Brilliant Blue. **B)** Band intensities of 3 ex-periments were measured using ImageJ software and normalized against CBB and relative to Col-0. Data points indicate individual experiments with points from the same experiment being marked with the same color (one-way ANOVA, Tukey’s post-hoc test; ns: not significant).

**Figure S4:**
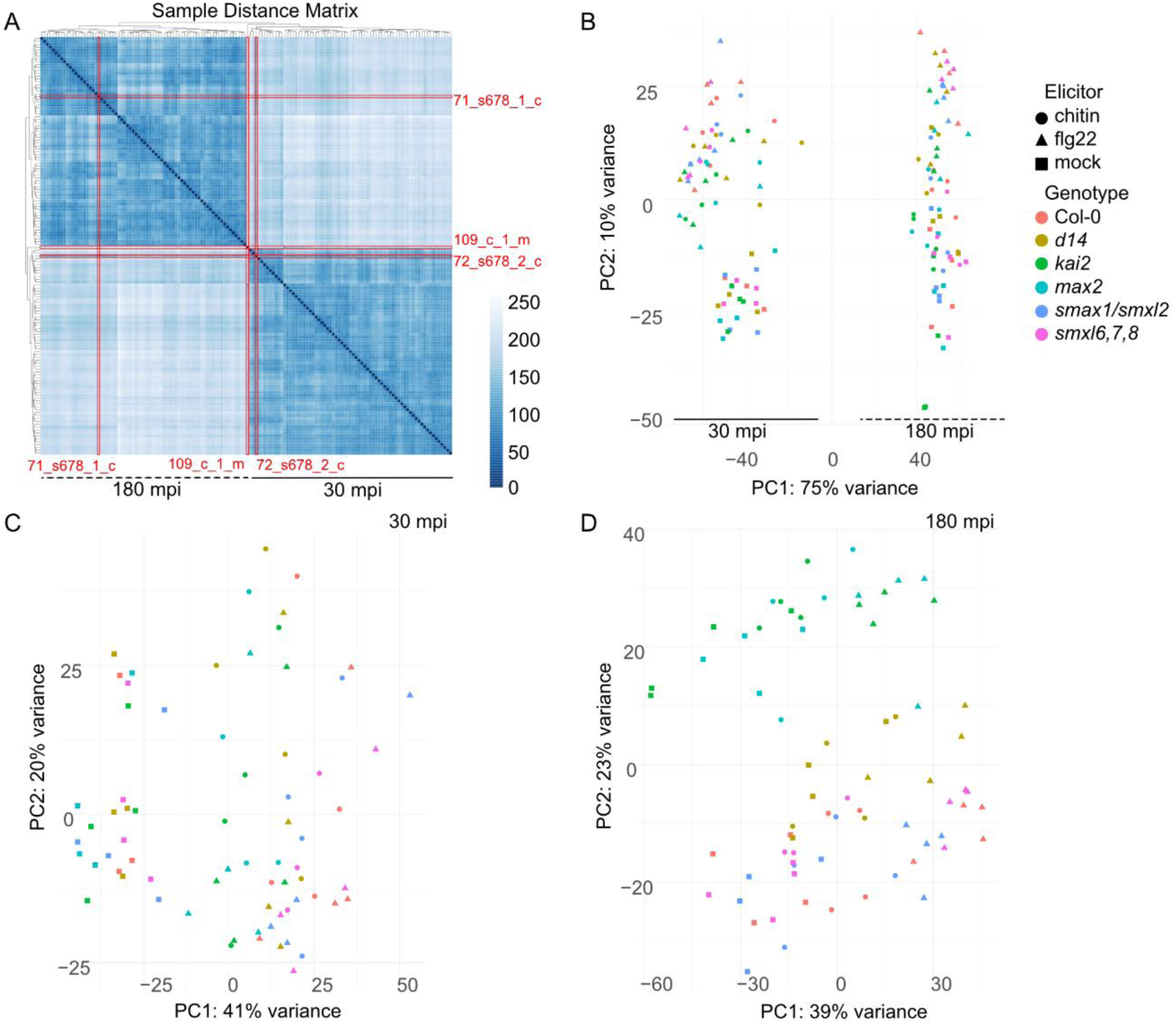
Hierarchical clustering and principal component analysis of global transcriptome profiles across genotypes, treatments, and timepoints. **A**) Hierarchical clustering of distance between samples in the RNA-seq dataset. Red rows and sample labels indicate samples removed prior to down-stream analysis due to separate clustering (109_c_1_m) or clustering with samples from the opposite timepoint (71_s678_1_c and 72_s678_2_c). **B**) Principal component analysis (PCA) of variance-stabi-lised gene expression values for all samples included in the dataset. Each point represents one biolog-ical replicate. The percentage of variance explained by each principal component is indicated on the respective axes. All points within -80 and 0 of PC1 are samples taken at 30 minutes post infiltration (mpi), and all points clustering upward of 0 of PC1 were harvested at 180 mpi. **C**) PCA of samples collected at 30 mpi using variance-stabilized expression values of all samples at 30 mpi. **D**) PCA of samples collected at using variance-stabilized expression values of all samples at 30 mpi. All panels display the first two principal components derived from the respective datasets.

**Fig. S5:**
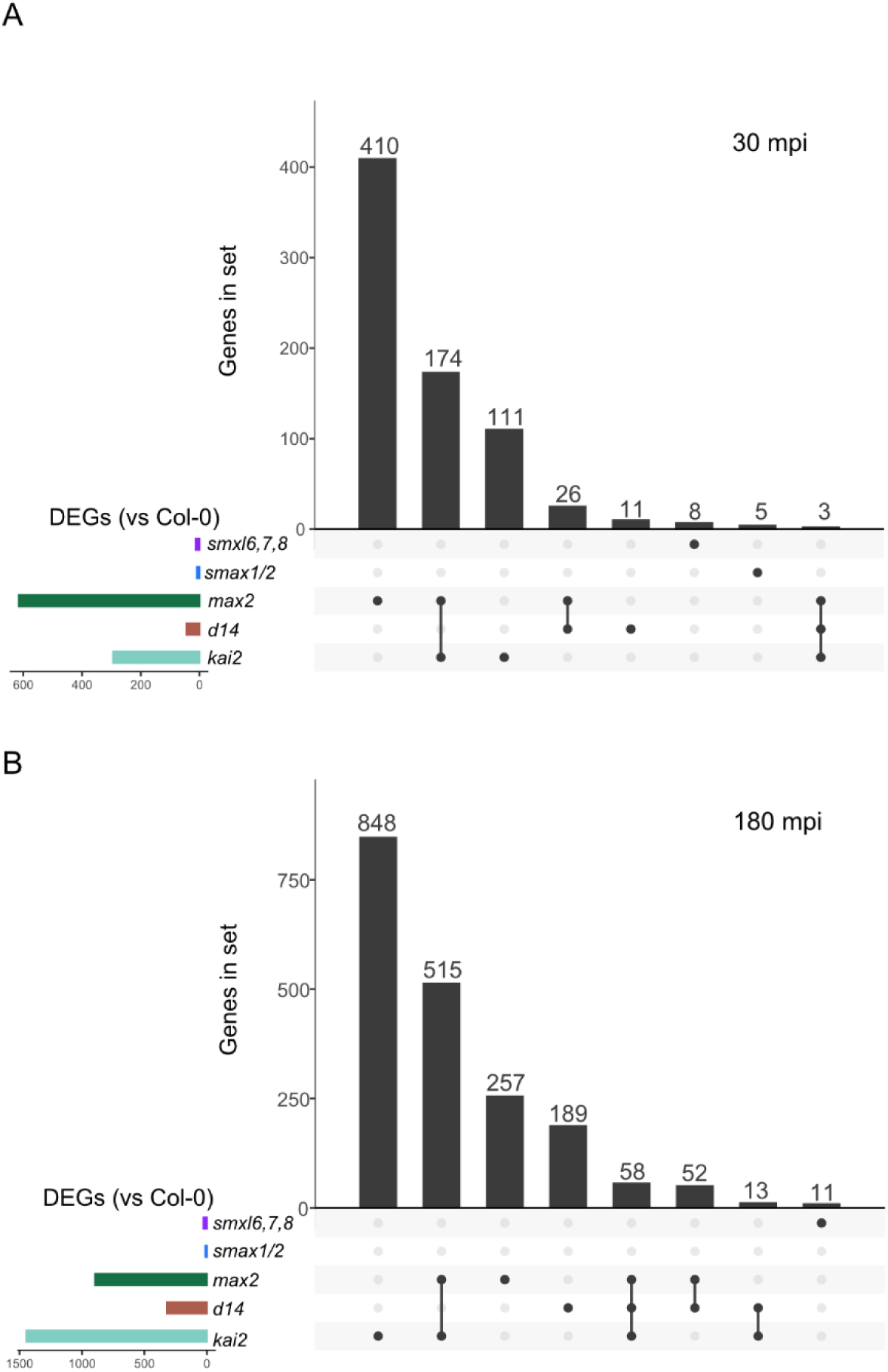
Upset plot of DEGs in mutant vs Col-0 leaves. Upset plots comparing the sets of differentially expressed genes (DEGs; padj < 0.05) in mutant vs Col-0 leaves under mock conditions at **A)** 30 minutes post infiltration (mpi) or **B)** 180 mpi. Vertical black bars and numbers correspond to the number of genes that are present in a set that can intersect across genotypes and connected dots below the bar indicate across which genotype vs Col-0 comparison genes are differentially expressed. Horizontal coloured bars indicate total amount of DEGs in genotype vs Col-0 comparison.

**Fig. S6:**
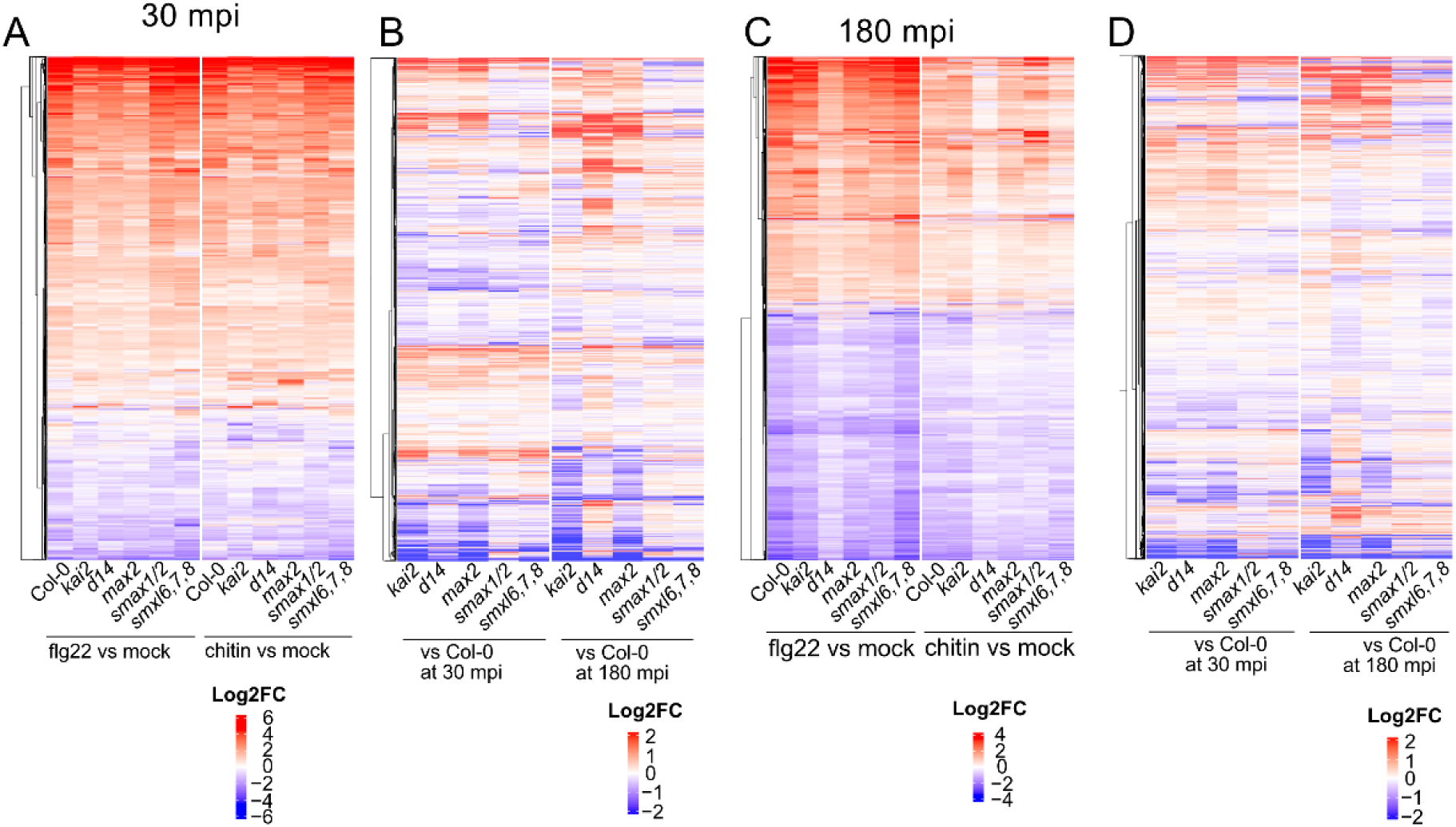
MAMP-induced and basal expression profile of all DEGs in leaves of Arabidopsis mu-tants treated with flg22 or chitin. Heatmaps showing Log_2_FC of all DEGs (DEGs; |Log_2_FC| > 1 & padj < 0.05) after flg22 or chitin infiltration at **A)** 30 minutes post infiltration (mpi) or **C)** 180 mpi and the basal expression across genotypes compared to Col-0 in **B)** and **D)** for the set of genes differentially regulated in **A)** or **C)** respectively.

**Fig. S7:**
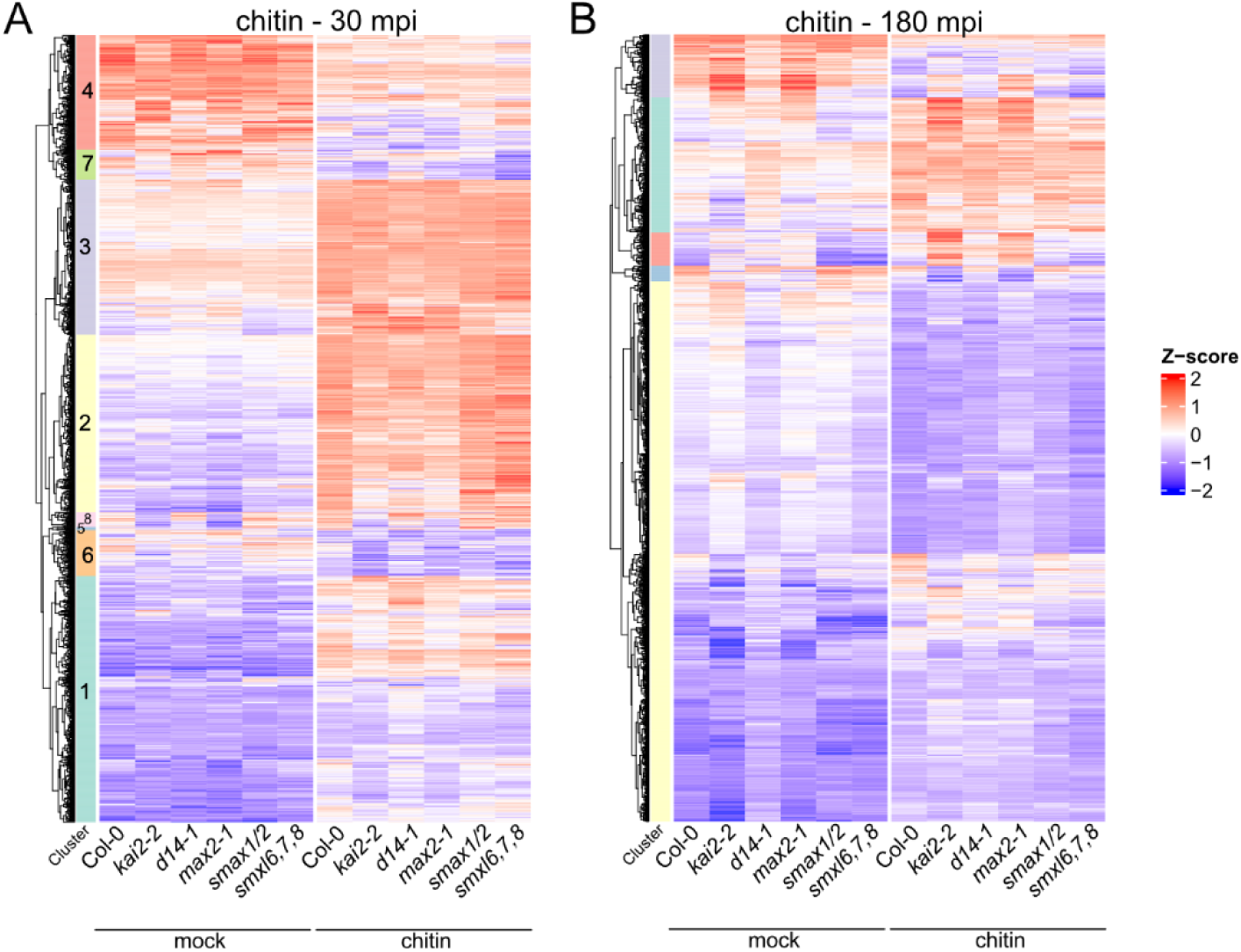
Chitin infiltration has a stronger effect on all genotypes at 30 compared to 180 minutes after infiltration. k-means clustered heatmaps of all DEGs (DEGs; |Log_2_FC| > 1 & padj < 0.05) with fold-change displayed as z-score in response to chitin treatment across genotypes at **A)** 30 minutes post infiltration (mpi) (1825 DEGs) and **B)** 180 mpi (1938 DEGs). Clusters were used for Gene Ontology enrichment in supplementary Figure S8.

**Fig. S8:**
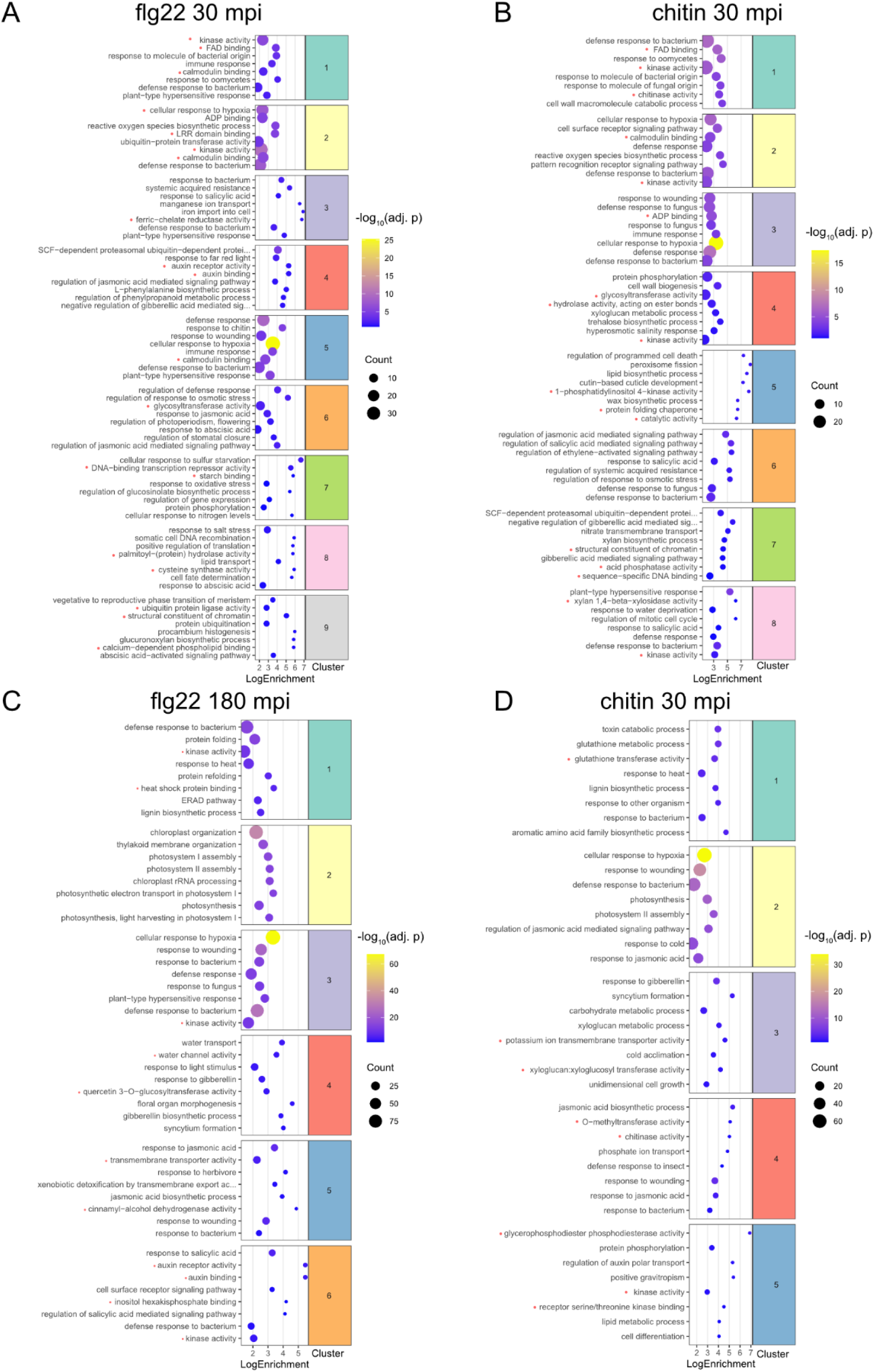
Gene Ontology term enrichment for gene clusters at 30 and 180 mpi after MAMP treat-ment. The 8 most significant Gene Ontology terms per cluster are shown for leaves treated with flg22 for **A)** 30 minutes post infiltration (mpi) or **B)** 180 mpi and with chitin for **C)** 30 mpi or **D)** 180 mpi. Colour of dot (blue to yellow) indicates enrichment significance (padj.) and size of dot indicates number of genes within the GO term. The x-axis indicates Log2Fold Enrichment. GO terms marked with a red dot indicate molecular function (MF). All other terms fall are categorized as biological process (BP).

**Fig. S9:**
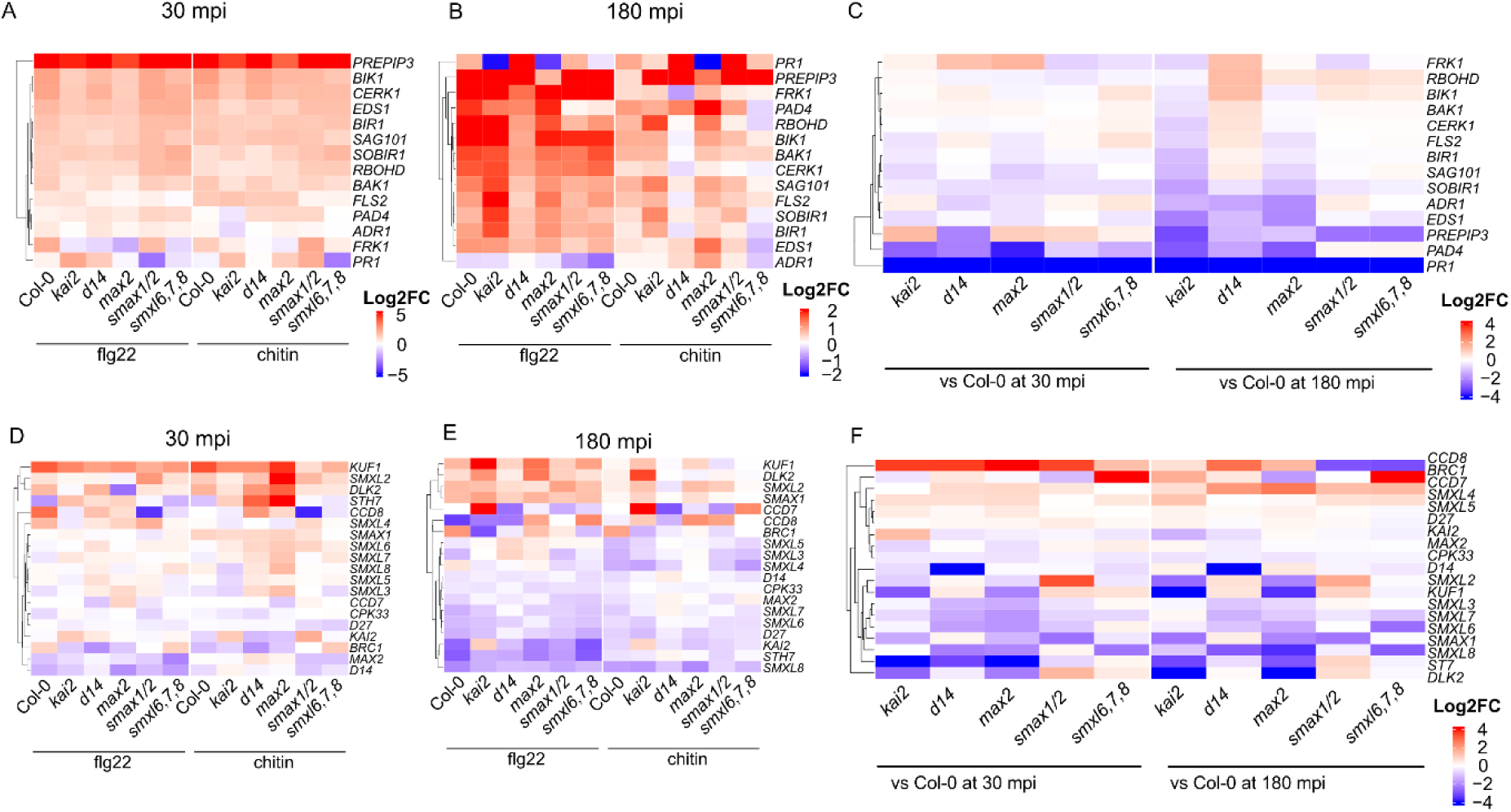
**Basal and MAMP-induced expression profile of immunity, karrikin and strigolactone sig-nalling-related genes.** Log_2_FC of gene expression in flg22- or chitin-treated vs. mock-treated leaves across genotypes for immunity-related (**A-C)** or KL and SL signalling/biosynthesis related genes (**D-F)** at **A, D)** 30 or **B, E)** 180 mpi. Log_2_FC of gene expression in mock-treated mutant vs. Col-0 leaves at 30 and 180 mpi for **C)** immunity-related or **F)** KL and SL signalling/biosynthesis genes.

**Fig. S10:**
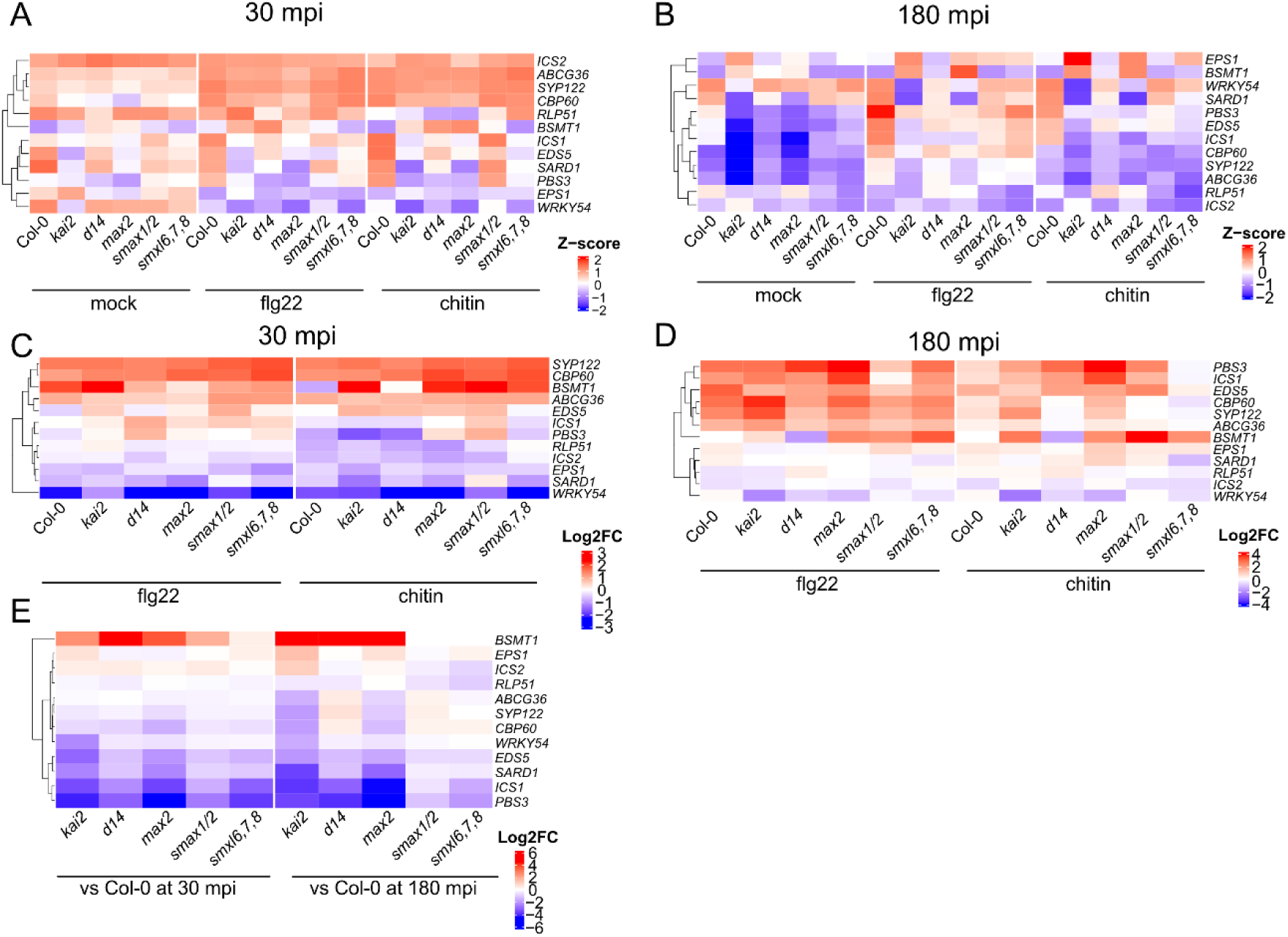
Basal and MAMP-induced expression profiles of SA biosynthesis and signalling genes. Median relative expression values (Z-values) **A-B)** or Log_2_FC **C-D)** of SA biosynthesis and signalling genes across leaves of all genotypes after mock-, flg22- or chitin-treatment at **A, C)** 30 or **B, D)** 180 mpi. **E)** Log_2_FC of SA biosynthesis and signalling genes in mutant vs. Col-0 leaves at 30 and 180 mpi after mock treatment.

**Fig. S11:**
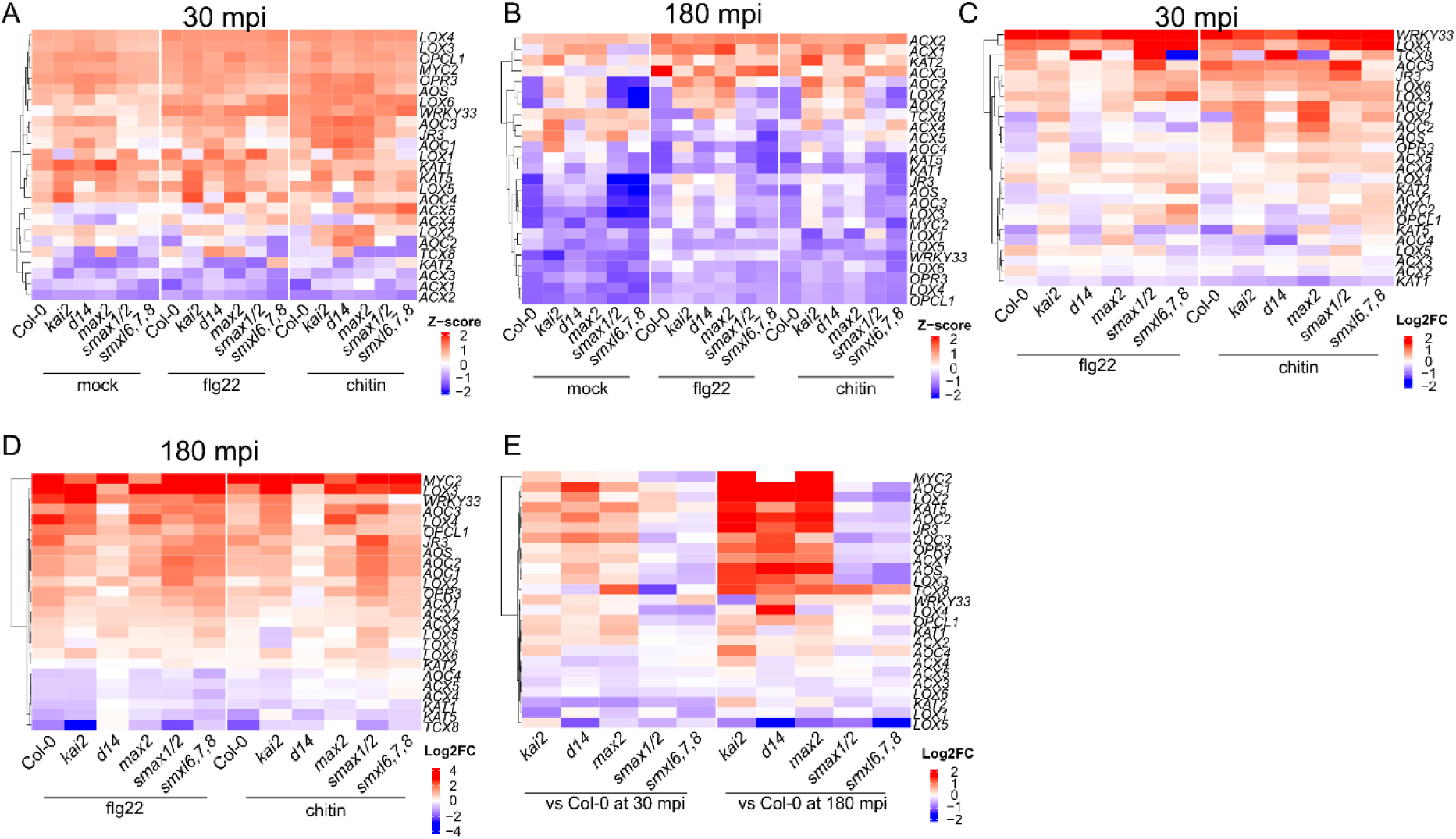
Basal and MAMP-induced expression profile of JA biosynthesis and signalling genes. Median relative expression values (Z-values) **A-B)** or Log_2_FC **C-D)** of JA biosynthesis and signalling genes across genotypes after mock-, flg22- or chitin-treatments at **A, C)** 30 or **B, D)** 180 mpi. **E)** Log_2_FC of JA biosynthesis and signalling genes in mutant vs Col-0 leaves at 30 and 180 mpi after mock treat-ment.

**Fig. S12:**
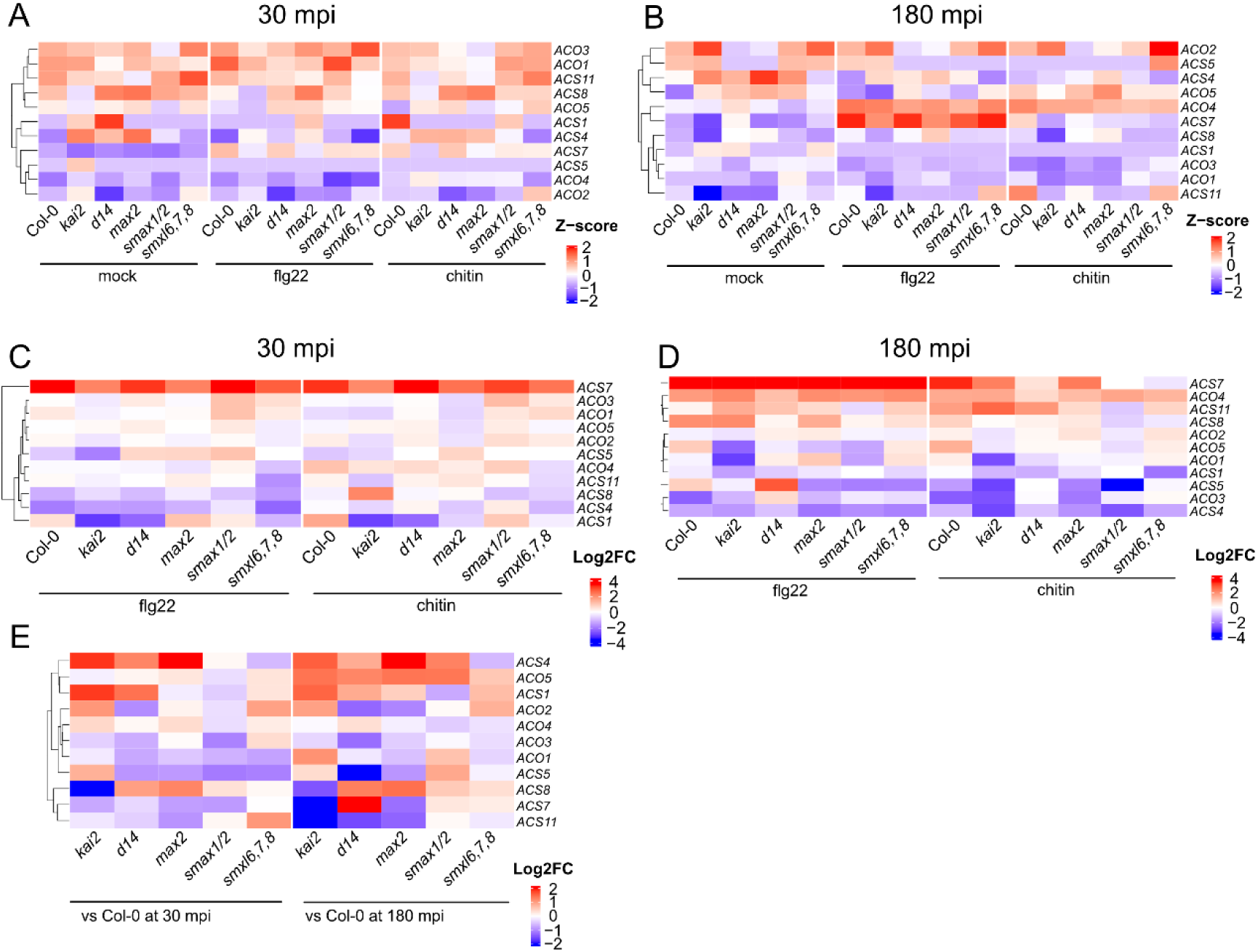
Basal and MAMP-induced expression profile of ethylene biosynthesis genes. Median relative expression values (Z-values) **A-B)** or Log_2_FC **C-D)** of ethylene biosynthesis genes across gen-otypes after mock-, flg22- or chitin-treatment at **A, C)** 30 or **B, D)** 180 mpi. **E)** Log_2_FC of ethylene bio-synthesis and signalling genes in mutant vs Col-0 leaves at 30 and 180 mpi after mock treatment.

**Fig. S13:**
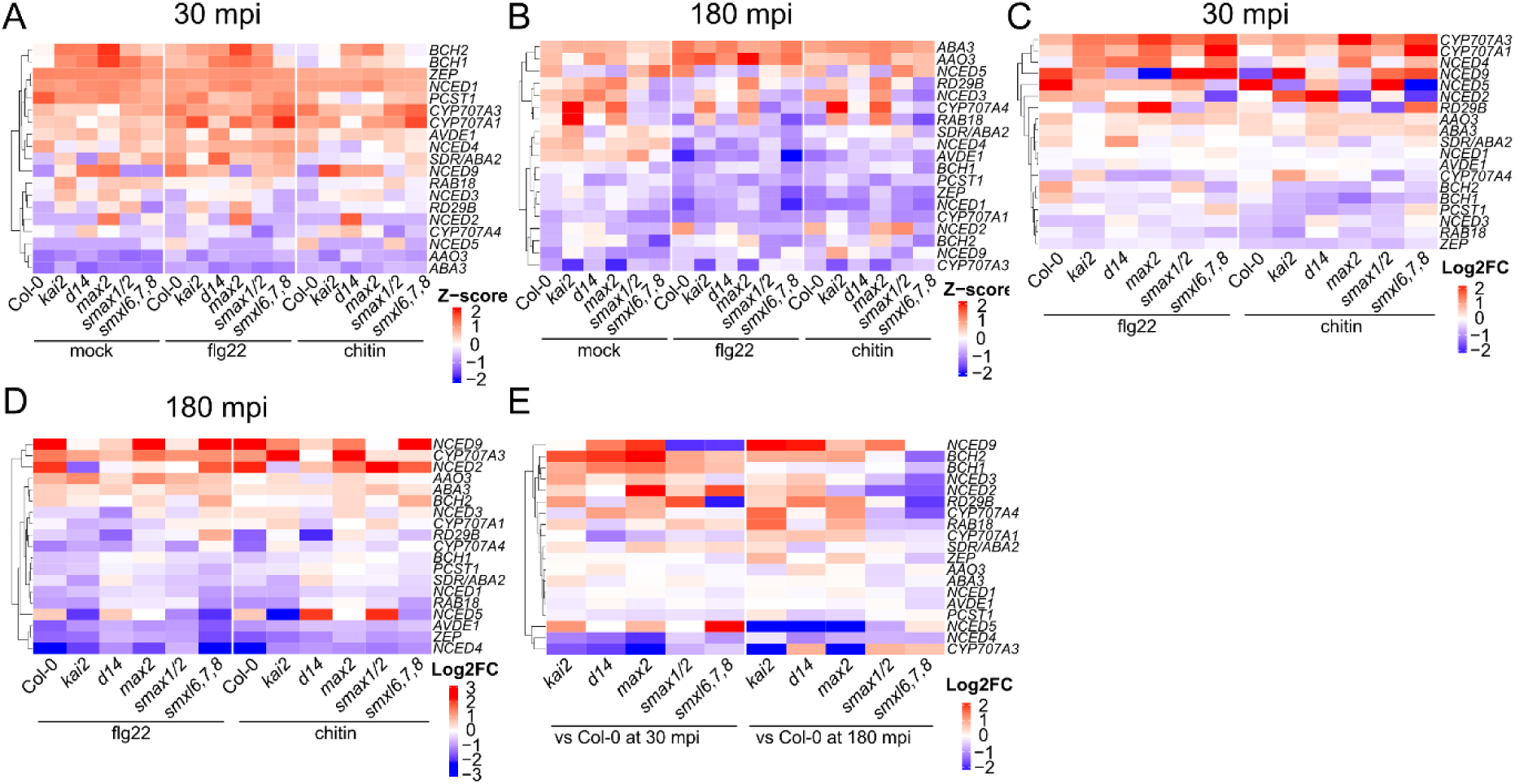
Basal and MAMP-induced expression profile of ABA biosynthesis genes. Median relative expression values (Z-values) **A-B)** or Log_2_FC **C-D)** of ABA biosynthesis genes across genotypes after mock-, flg22- or chitin-treatment at **A, C)** 30 or **B, D)** 180 mpi. **E)** Log_2_FC of ABA biosynthesis genes in mutant vs Col-0 leaves at 30 and 180 mpi after mock treatment.

**Table 1:** Plant lines used in this study.

| Plant line | Reference |
| --- | --- |
| ***Arabidopsis thaliana*** |  |
| *kai2-1* | (100) |
| *kai2-2* | (100) |
| *max2-1* | (101) |
| *max2-2* | (101) |
| *d14-1* | (18) |
| *smax1-2 smxl2-1* | (19) |
| *smxl6 smxl7 smxl8* | (37) |
| ***L. japonicus*** |  |
| *kai2a-1 kai2b-1* | (13) |
| *d14-1* | (13) |
| *max2-3* | (13) |
| *smax1-3* | (13) |
| ***N. benthamiana*** |  |
| *kai2a kai2b kai2c kai2d* | (49) |

## References

1. Doughari J. An overview of plant immunity. J Plant Pathol Microbiol. 2015;6(322):1–11. doi:10.4172/2157-7471.1000322

2. Bürger M, Chory J. Stressed out about hormones: How plants orchestrate immunity. Cell Host Microbe. 2019; 26(2):163–72. doi:10.1016/j.chom.2019.07.006 PMID: 31415749.

3. Shafqat A, Abbas S, Ambreen M, Siddiqa Bhatti A, Kausar H, Gull T. Exploring the vital role of phytohormones and plant growth regulators in orchestrating plant immunity. Physiol Mol Plant Pathol. 2024; 133:102359. doi:10.1016/j.pmpp.2024.102359

4. Varshney K, Gutjahr C. KAI2 can do: karrikin receptor function in plant development and response to abiotic and biotic factors. Plant Cell Physiol. 2023; 64(9):984–95. doi:10.1093/pcp/pcad077 PMID: 37548562.

5. Flematti GR, Ghisalberti EL, Dixon KW, Trengove RD. A compound from smoke that promotes seed germination. Science; 305(5686): 977. doi:10.1126/science.1099944 PMID: 15247439.

6. Nelson DC, Flematti GR, Riseborough JA, Ghisalberti EL, Dixon KW, Smitha SM. Karrikins enhance light responses during germination and seedling development in *Arabidopsis thaliana*. Proc Natl Acad Sci U S A. 2010; 107(15): 7095–100. doi:10.1073/pnas.0911635107 PMID: 20351290.

7. Waters MT, Nelson DC. Karrikin perception and signalling. New Phytol. 2023; 237(5):1525–41. doi:10.1111/nph.18598 PMID: 36333982.

8. Nelson DC, Scaffidi A, Dun EA, Waters MT, Flematti GR, Dixon KW, et al. F-box protein MAX2 has dual roles in karrikin and strigolactone signaling in *Arabidopsis thaliana*. Proc Natl Acad Sci U S A. 2011; 108(21): 8897–902. doi:10.1073/pnas.1100987108 PMID: 21555559.

9. Li W, Nguyen KH, Chu HD, Ha C Van, Watanabe Y, Osakabe Y, et al. The karrikin receptor KAI2 promotes drought resistance in *Arabidopsis thaliana*. PLoS Genet. 2017; 13(11):e1007076. doi:10.1371/journal.pgen.1007076 PMID: 29131815.

10. Feng Z, Liang X, Tian H, Watanabe Y, Nguyen KH, Tran CD, et al. SUPPRESSOR of MAX2 1 (SMAX1) and SMAX1-LIKE2 (SMXL2) negatively regulate drought resistance in *Arabidopsis thaliana*. Plant Cell Physiol. 2023;63(12):1900–13. doi:10.1093/pcp/pcac080 PMID: 35681253.

11. Swarbreck SM, Guerringue Y, Matthus E, Jamieson FJC, Davies JM. Impairment in karrikin but not strigolactone sensing enhances root skewing in *Arabidopsis thaliana*. Plant J. 2019; 98(4): 607–21. doi:10.1111/tpj.14233 PMID: 30659713.

12. Villaécija-Aguilar JA, Hamon-Josse M, Carbonnel S, Kretschmar A, Schmidt C, Dawid C, et al. SMAX1/SMXL2 regulate root and root hair development downstream of KAI2-mediated signal-ling in Arabidopsis. PLoS Genet. 2019; 15(8): e1008327. doi:10.1371/journal.pgen.1008327 PMID: 31465451.

13. Carbonnel S, Das D, Varshney K, Kolodziej MC, Villaécija-Aguilar JA, Gutjahr C. The karrikin signaling regulator SMAX1 controls *Lotus japonicus* root and root hair development by sup-pressing ethylene biosynthesis. Proc Natl Acad Sci U S A. 2020; 117(35): 21757–65. doi:10.1073/pnas.2006111117 PMID: 32817510.

14. Waters MT, Scaffidi A, Moulin SLY, Sun YK, Flematti GR, Smith SM. A *Selaginella moellen-dorffii* ortholog of KARRIKIN INSENSITIVE2 functions in Arabidopsis development but cannot mediate responses to karrikins or strigolactones. Plant Cell. 2015; 27(7):1925–44. doi:10.1105/tpc.15.00146 PMID: 26175507.

15. Sepulveda C, Guzmán MA, Li Q, Antonio Villaécija-Aguilar J, Martinez SE, Kamran M, et al. KARRIKIN UP-REGULATED F-BOX 1 (KUF1) imposes negative feedback regulation of karri-kin and KAI2 ligand metabolism in *Arabidopsis thaliana*. Proc Natl Acad Sci U S A. 2022; 119(11):e2112820119. doi:10.1073/pnas.2112820119 PMID: 35254909.

16. Conn CE, Nelson DC. Evidence that KARRIKIN-INSENSITIVE2 (KAI2) receptors may perceive an unknown signal that is not karrikin or strigolactone. Front Plant Sci. 2016; 6:1219. doi:10.3389/fpls.2015.01219 PMID: 26779242.

17. Sun YK, Flematti GR, Smith SM, Waters MT. Reporter gene-facilitated detection of compounds in Arabidopsis leaf extracts that activate the karrikin signaling pathway. Front Plant Sci. 2016; 7:1799. doi:10.3389/fpls.2016.01799 PMID: 27994609.

18. Waters MT, Nelson DC, Scaffidi A, Flematti GR, Sun YK, Dixon KW, et al. Specialisation within the DWARF14 protein family confers distinct responses to karrikins and strigolactones in Ara-bidopsis. Development. 2012; 139(7): 1285–95. doi:10.1242/dev.074567 PMID: 22357928.

19. Stanga JP, Morffy N, Nelson DC. Functional redundancy in the control of seedling growth by the karrikin signaling pathway. Planta. 2016; 243(6):1397–406. doi:10.1007/s00425-015-2458-2 PMID: 26754282.

20. Stanga JP, Smith SM, Briggs WR, Nelson DC. SUPPRESSOR OF MORE AXILLARY GROWTH2 1 controls seed germination and seedling development in Arabidopsis. Plant Phys-iol. 2013; 163(1): 318–30. doi:10.1104/pp.113.221259 PMID: 23893171.

21. Walker CH, Siu-Ting K, Taylor A, O’Connell MJ, Bennett T. Strigolactone synthesis is ancestral in land plants, but canonical strigolactone signalling is a flowering plant innovation. BMC Biol. 2019; 17(1): 70. doi:10.1186/s12915-019-0689-6 PMID: 31488154.

22. Vancea AI, Huntington B, Steinchen W, Savva CG, Shahul Hameed UF, Arold ST. Mechanism of cooperative strigolactone perception by the MAX2 ubiquitin ligase–receptor–substrate com-plex. Nat Commun. 2025; 16(1): 10291. doi:10.1038/s41467-025-65205-0 PMID: 41271672.

23. Khosla A, Morffy N, Li Q, Faure L, Chang SH, Yao J, et al. Structure–Function analysis of SMAX1 reveals domains that mediate its karrikin-induced proteolysis and interaction with the receptor KAI2. Plant Cell. 2020; 32(8): 2639–59. doi:10.1105/tpc.19.00752 PMID: 32434855.

24. Wang L, Xu Q, Yu H, Ma H, Li X, Yang J, et al. Strigolactone and karrikin signaling pathways elicit ubiquitination and proteolysis of SMXL2 to regulate hypocotyl elongation in Arabidopsis. Plant Cell. 2020; 32(7):2251–70. doi:10.1105/tpc.20.00140 PMID: 32358074.

25. Li Q, Martín-Fontecha ES, Khosla A, White ARF, Chang S, Cubas P, et al. The strigolactone receptor D14 targets SMAX1 for degradation in response to GR24 treatment and osmotic stress. Plant Commun. 2022; 3(2): 100303. doi:10.1016/j.xplc.2022.100303 PMID: 35529949.

26. Challis RJ, Hepworth J, Mouchel C, Waites R, Leyser O. A role for More Axillary Growth1 (MAX1) in evolutionary diversity in strigolactone signaling upstream of MAX2. Plant Physiol. 2013; 161(4): 1885–902. doi:10.1104/pp.112.211383 PMID: 23424248.

27. Kapulnik Y, Delaux PM, Resnick N, Mayzlish-Gati E, Wininger S, Bhattacharya C, et al. Strigo-lactones affect lateral root formation and root-hair elongation in Arabidopsis. Planta. 2011; 233(1):209–16. doi:10.1007/s00425-010-1310-y PMID: 21080198.

28. Kapulnik Y, Resnick N, Mayzlish-Gati E, Kaplan Y, Wininger S, Hershenhorn J, et al. Strigolac-tones interact with ethylene and auxin in regulating root-hair elongation in Arabidopsis. J Exp Bot. 2011; 62(8): 2915–24. doi:10.1093/jxb/erq464 PMID: 21307387.

29. Umehara M, Hanada A, Yoshida S, Akiyama K, Arite T, Takeda-Kamiya N, et al. Inhibition of shoot branching by new terpenoid plant hormones. Nature. 2008; 455(7210): 195–200. doi:10.1038/nature07272 PMID: 18690207.

30. Gomez-Roldan V, Fermas S, Brewer PB, Puech-Pagès V, Dun EA, Pillot JP, et al. Strigolac-tone inhibition of shoot branching. Nature. 2008; 455(7210): 189–94. doi:10.1038/nature07271 PMID: 18690209.

31. Ueda H, Kusaba M. Strigolactone regulates leaf senescence in concert with ethylene in Ara-bidopsis. Plant Physiol. 2015; 169(1): 138–47. doi:10.1104/pp.15.00325 PMID: 25979917.

32. Agusti J, Herold S, Schwarz M, Sanchez P, Ljung K, Dun EA, et al. Strigolactone signaling is required for auxin-dependent stimulation of secondary growth in plants. Proc Natl Acad Sci U S A. 2011; 108(50): 20242–7. doi:10.1073/pnas.1111902108 PMID: 22123958.

33. Hamiaux C, Drummond RSM, Janssen BJ, Ledger SE, Cooney JM, Newcomb RD, et al. DAD2 is an α/β hydrolase likely to be involved in the perception of the plant branching hormone, strigolactone. Curr Biol. 2012; 22(21): 2032–6. doi:10.1016/j.cub.2012.08.007 PMID: 22959345.

34. Yao R, Ming Z, Yan L, Li S, Wang F, Ma S, et al. DWARF14 is a non-canonical hormone re-ceptor for strigolactone. Nature. 2016; 536(7617): 469–73. doi:10.1038/nature19073 PMID: 27479325.

35. Bythell-Douglas R, Rothfels CJ, Stevenson DWD, Graham SW, Wong GKS, Nelson DC, et al. Evolution of strigolactone receptors by gradual neo-functionalization of KAI2 paralogues. BMC Biol. 2017; 15(1):52. doi:10.1186/s12915-017-0397-z PMID: 28662667.

36. Delaux PM, Xie X, Timme RE, Puech-Pages V, Dunand C, Lecompte E, et al. Origin of strigo-lactones in the green lineage. New Phytol. 2012; 195(4): 857–71. doi:10.1111/j.1469-8137.2012.04209.x PMID: 22738134.

37. Soundappan I, Bennett T, Morffy N, Liang Y, Stanga JP, Abbas A, et al. SMAX1-LIKE/D53 family members enable distinct MAX2-dependent responses to strigolactones and karrikins in Arabidopsis. Plant Cell. 2015; 27(11): 3143–59. doi:10.1105/tpc.15.00562 PMID: 26546447.

38. Wang L, Wang B, Jiang L, Liu X, Li X, Lu Z, et al. Strigolactone signaling in Arabidopsis regu-lates shoot development by targeting D53-like SMXL repressor proteins for ubiquitination and degradation. Plant Cell. 2015; 27(11): 3128–42. doi:10.1105/tpc.15.00605 PMID: 26546446.

39. Zhou F, Lin Q, Zhu L, Ren Y, Zhou K, Shabek N, et al. D14-SCF^D3^ -dependent degradation of D53 regulates strigolactone signalling. Nature. 2013; 504(7480): 406–10. doi:10.1038/na-ture12878 PMID: 24336215.

40. Jiang L, Liu X, Xiong G, Liu H, Chen F, Wang L, et al. DWARF 53 acts as a repressor of strigo-lactone signalling in rice. Nature. 2013; 504(7480): 401–5. doi:10.1038/nature12870 PMID: 24336200.

41. Chang SH, George WJ, Nelson DC. An N-terminal domain specifies developmental control by the SMAX1-LIKE family of transcriptional regulators in *Arabidopsis thaliana*. Proc Natl Acad Sci U S A. 2025; 122(24): e2412793122. doi:10.1073/pnas.2412793122 PMID: 40493196.

42. Ma H, Duan J, Ke J, He Y, Gu X, Xu TH, et al. A D53 repression motif induces oligomerization of TOPLESS corepressors and promotes assembly of a corepressor-nucleosome complex. Sci Adv. 2017; 3(6). doi:10.1126/sciadv.1601217 PMID: 28630893.

43. Wang L, Wang B, Yu H, Guo H, Lin T, Kou L, et al. Transcriptional regulation of strigolactone signalling in Arabidopsis. Nature. 2020; 583(7815):277–81. doi:10.1038/s41586-020-2382-x PMID: 32528176.

44. Das D, Varshney K, Ogawa S, Torabi S, Hüttl R, Nelson DC, et al. Ethylene promotes SMAX1 accumulation to inhibit arbuscular mycorrhiza symbiosis. Nat Commun. 2025; 16(1):2025. doi:10.1038/s41467-025-57222-w PMID: 40016206.

45. Gutjahr C, Gobbato E, Choi J, Riemann M, Johnston MG, Summers W, et al. Rice perception of symbiotic arbuscular mycorrhizal fungi requires the karrikin receptor complex. Science. 2015; 350(6267): 1521–4. doi:10.1126/science.aac9715 PMID: 26680197.

46. Choi J, Lee T, Cho J, Servante EK, Pucker B, Summers W, et al. The negative regulator SMAX1 controls mycorrhizal symbiosis and strigolactone biosynthesis in rice. Nat Commun. 2020; 11: 2114. doi:10.1038/s41467-020-16021-1 PMID: 32355217.

47. Meng Y, Varshney K, Incze N, Badics E, Kamran M, Davies SF, et al. KARRIKIN INSENSI-TIVE2 regulates leaf development, root system architecture and arbuscular-mycorrhizal symbi-osis in *Brachypodium distachyon*. Plant J. 2022; 109(6): 1559–74. doi:10.1111/tpj.15651 PMID: 34953105.

48. Li XR, Sun J, Albinsky D, Zarrabian D, Hull R, Lee T, et al. Nutrient regulation of lipochitooligo-saccharide recognition in plants via NSP1 and NSP2. Nat Commun. 2022; 13(1): 6421. doi:10.1038/s41467-022-33908-3 PMID: 36307431.

49. Buhrmann K, Torabi S, Carbonnel S, Varshney K, Chapman P, Fenn A, et al. Unequal require-ment of KAI2 for AM symbiosis across angiosperms. bioRxiv. 2026. doi:10.64898/2026.05.03.722480

50. Rehman N ur, Ali M, Ahmad MZ, Liang G, Zhao J Strigolactones promote rhizobia interaction and increase nodulation in soybean (*Glycine max*). Microb Pathog. 2018; 114:420–30. doi:10.1016/j.micpath.2017.11.049 PMID: 29191709.

51. Akiyama K, Matsuzaki KI, Hayashi H. Plant sesquiterpenes induce hyphal branching in arbus-cular mycorrhizal fungi. Nature. 2005; 435(7043): 824–7. doi:10.1038/nature03608 PMID: 15944706.

52. Foo E, Yoneyama K, Hugill CJ, Quittenden LJ, Reid JB. Strigolactones and the regulation of pea symbioses in response to nitrate and phosphate deficiency. Mol Plant. 2013; 6(1): 76–87. doi:10.1093/mp/sss115 PMID: 23066094.

53. Buhrman K, Gutjahr C. Regulation of arbuscular mycorrhiza development by environmental stimuli: Many roads lead to strigolactones. PLoS Pathog. 2025; 21:e1013555. doi:10.1371/jour-nal.ppat.1013555 PMID: 41060984.

54. Zhang C, He J, Dai H, Wang G, Zhang X, Wang C, et al. Discriminating symbiosis and immun-ity signals by receptor competition in rice. Proc Natl Acad Sci U S A. 2021; 118:20237–38118. doi:10.1073/pnas.2023738118 PMID: 33853950.

55. Tan X, Wang D, Zhang X, Zheng S, Jia X, Liu H, et al. A pair of LysM receptors mediates sym-biosis and immunity discrimination in Marchantia. Cell. 2025; 188(5): 1330–1348.e27. doi:10.1016/j.cell.2024.12.024 PMID: 39855200.

56. Wang D, Jin R, Shi X, Guo H, Tan X, Zhao A, et al. A kinase mediator of rhizobial symbiosis and immunity in Medicago. Nature. 2025; 643(8072): 768–75. doi:10.1038/s41586-025-09057-0 PMID: 40328313.

57. Zhang J, Sun J, Chiu CH, Landry D, Li K, Wen J, et al. A receptor required for chitin perception facilitates arbuscular mycorrhizal associations and distinguishes root symbiosis from immunity. Curr Biol. 2024; 34(8): 1705–1717.e6. doi:10.1016/j.cub.2024.03.015 PMID: 38574729.

58. Rey T, Nars A, Bonhomme M, Bottin A, Huguet S, Balzergue S, et al. NFP, a LysM protein controlling Nod factor perception, also intervenes in *Medicago truncatula* resistance to patho-gens. New Phytol. 2013; 198(3): 875–86. doi:10.1111/nph.12198 PMID: 23432463.

59. Tsitsikli M, Simonsen B, Luu TB, Larsen MM, Andersen CG, Gysel K, et al. Two residues re-program immunity receptors for nitrogen-fixing symbiosis. Nature. 2025; 648(8093): 443–50. doi:10.1038/s41586-025-09696-3 PMID: 41193803.

60. Bozsoki Z, Gysel K, Hansen SB, Lironi D, Krönauer C, Feng F, et al. Ligand-recognizing motifs in plant LysM receptors are major determinants of specificity. Science. 2020; 369(6504): 663–70. doi:10.1126/science.abb3377 PMID: 32764065.

61. Jones JDG, Staskawicz BJ, Dangl JL. The plant immune system: From discovery to deploy-ment. Cell. 2024; 187(9): 2095–116. doi:10.1016/j.cell.2024.03.045 PMID: 38670067.

62. DeFalco TA, Zipfel C. Molecular mechanisms of early plant pattern-triggered immune signaling. Mol Cell. 2021; 81(17): 3449–67. doi:10.1016/j.molcel.2021.07.029 PMID: 34403694.

63. Bigeard J, Colcombet J, Hirt H. Signaling mechanisms in pattern-triggered immunity (PTI). Mol Plant. 2015; 8(4): 521–39. doi:10.1016/j.molp.2014.12.022 PMID: 25744358.

64. Ngou BPM, Ding P, Jones JDG. Thirty years of resistance: Zig-zag through the plant immune system. Plant Cell. 2022; 34(5): 1447–78. doi:10.1093/plcell/koac041 PMID: 35167697.

65. Stes E, Depuydt S, De Keyser A, Matthys C, Audenaert K, Yoneyama K, et al. Strigolactones as an auxiliary hormonal defence mechanism against leafy gall syndrome in *Arabidopsis thali-ana*. J Exp Bot. 2015; 66(16): 5123–34. doi:10.1093/jxb/erv309 PMID: 26136271.

66. Piisilä M, Keceli MA, Brader G, Jakobson L, Jöesaar I, Sipari N, et al. The F-box protein MAX2 contributes to resistance to bacterial phytopathogens in *Arabidopsis thaliana*. BMC Plant Biol. 2015; 15(1): 53. doi:10.1186/s12870-015-0434-4 PMID: 25849639.

67. Kusajima M, Fujita M, Soudthedlath K, Nakamura H, Yoneyama K, Nomura T, et al. Strigolac-tones modulate salicylic acid-mediated disease resistance in *Arabidopsis thaliana*. Int J Mol Sci. 2022; 23(9): 5246. doi:10.3390/ijms23095246 PMID: 35563637.

68. Foo E, Blake SN, Fisher BJ, Smith JA, Reid JB. The role of strigolactones during plant interac-tions with the pathogenic fungus *Fusarium oxysporum*. Planta. 2016; 243(6): 1387–96. doi:10.1007/s00425-015-2449-3 PMID: 26725046.

69. Nasir F, Tian L, Shi S, Chang C, Ma L, Gao Y, et al. Strigolactones positively regulate defense against *Magnaporthe oryzae* in rice (*Oryza sativa*). Plant Physiol Biochem. 2019; 142: 106–16. doi:10.1016/j.plaphy.2019.06.028 PMID: 31279135.

70. Felix G, Duran JD, Volko S, Boller T. Plants have a sensitive perception system for the most conserved domain of bacterial flagellin. Plant J. 1999; 18(3): 265–76. doi:10.1046/j.1365-313X.1999.00265.x PMID: 10377992.

71. Gómez-Gómez L, Boller T. FLS2: An LRR receptor-like kinase involved in the perception of the bacterial elicitor flagellin in Arabidopsis. Mol Cell. 2000; 5(6): 1003–11. doi:10.1016/S1097-2765(00)80265-8 PMID: 10911994.

72. Miya A, Albert P, Shinya T, Desaki Y, Ichimura K, Shirasu K, et al. CERK1, a LysM receptor kinase, is essential for chitin elicitor signaling in Arabidopsis. Proc Natl Acad Sci U S A. 2007; 104(49): 19613–8. doi:10.1073/pnas.0705147104 PMID: 18042724.

73. Angel TM, Dangl JL, G Jones JD. Arabidopsis gp91^phox^ homologues AtrbohD and AtrbohF are required for accumulation of reactive oxygen intermediates in the plant defense response. Proc Natl Acad Sci U S A. 2002; 99(1): 517–22. doi:10.1073/pnas.012452499 PMID: 11756663.

74. Zipfel C, Robatzek S, Navarro L, Oakeley EJ, Jones JDG, Felix G, et al. Bacterial disease re-sistance in Arabidopsis through flagellin perception. Nature. 2004; 428: 764–7. doi:10.1038/na-ture02485 PMID: 15085136.

75. Bjornson M, Pimprikar P, Nürnberger T, Zipfel C. The transcriptional landscape of *Arabidopsis thaliana* pattern-triggered immunity. Nat Plants. 2021; 7(5): 579–86. doi:10.1038/s41477-021-00874-5 PMID: 33723429.

76. Zheng X, Liu F, Yang X, Li W, Chen S, Yue X, et al. The MAX2-KAI2 module promotes salicylic acid-mediated immune responses in Arabidopsis. J Integr Plant Biol. 2023; 65(6): 1566–84. doi:10.1111/jipb.13463 PMID: 36738234.

77. Kusajima M, Fujita M, Takahashi I, Mori T, Tanaka T, Nakamura H, et al. Enhancement of sys-temic acquired resistance in rice by F-box protein D3-mediated strigolactone/karrikin signaling. Sci Rep. 2025; 15(1): 23875. doi:10.1038/s41598-025-06984-w PMID: 40615560.

78. Salem MA, Jamil M, Wang JY, Berqdar L, Liew KX, Paramita A, et al. Disruption of the karrikin receptor DWARF 14 LIKE (D14L) gene leads to distinct effects on root and shoot growth, and reprogramming of central metabolism in rice. J Exp Bot. 2025; 76(14): 4114–28. doi:10.1093/jxb/eraf201 PMID: 40357891.

79. Simons JL, Napoli CA, Janssen BJ, Plummer KM, Snowden KC. Analysis of the DECREASED APICAL DOMINANCE genes of petunia in the control of axillary branching. Plant Physiol. 2007; 143(2): 697–706. doi:10.1104/pp.106.087957 PMID: 17158589.

80. Bozsoki Z, Cheng J, Feng F, Gysel K, Vinther M, Andersen KR, et al. Receptor-mediated chitin perception in legume roots is functionally separable from Nod factor perception. Proc Natl Acad Sci U S A. 2017; 114(38): E8118–27. doi:10.1073/pnas.1706795114 PMID: 28874587.

81. Lian Y, Lian C, Wang L, Li Z, Yuan G, Xuan L, et al. SUPPRESSOR OF MAX2 LIKE 6, 7, and 8 interact with DDB1 BINDING WD REPEAT DOMAIN HYPERSENSITIVE TO ABA DEFI-CIENT 1 to regulate the drought tolerance and target SUCROSE NONFERMENTING 1 RE-LATED PROTEIN KINASE 2.3 to abscisic acid response in Arabidopsis. Biomolecules. 2023; 13(9): 1406. doi:10.3390/biom13091406 PMID: 37759806.

82. Li N, Han X, Feng D, Yuan D, Huang LJ. Signaling crosstalk between salicylic acid and eth-ylene/jasmonate in plant defense: Do we understand what they are whispering? Int J Mol Sci. 2019; 20(3): 671. doi:10.3390/ijms20030671 PMID: 30720746.

83. Macho AP, Boutrot F, Rathjen JP, Zipfel C. ASPARTATE OXIDASE plays an important role in Arabidopsis stomatal immunity. Plant Physiol. 2012; 159(4): 1845–56. doi:10.1104/pp.112.199810 PMID: 22730426.

84. Fichman Y, Zandalinas SI, Peck S, Luan S, Mittler R. HPCA1 is required for systemic reactive oxygen species and calcium cell-to-cell signaling and plant acclimation to stress. Plant Cell. 2022; 34(11): 4453–71. doi:10.1093/plcell/koac241 PMID: 35929088.

85. Dubiella U, Seybold H, Durian G, Komander E, Lassig R, Witte CP, et al. Calcium-dependent protein kinase/NADPH oxidase activation circuit is required for rapid defense signal propaga-tion. Proc Natl Acad Sci U S A. 2013; 110(21): 8744–9. doi:10.1073/pnas.1221294110 PMID: 23650383.

86. Köster P, He G, Liu C, Dong Q, Hake K, Schmitz-Thom I, et al. A bi-kinase module sensitizes and potentiates plant immune signaling. Sci Adv. 2025; 11(4): eadt9804. doi:10.1126/sci-adv.adt9804 PMID: 39854470.

87. Stegmann M, Monaghan J, Smakowska-Luzan E, Rovenich H, Lehner A, Holton N, et al. The receptor kinase FER is a RALF-regulated scaffold controlling plant immune signaling. Science. 2017; 355(6322): 287–9. doi:10.1126/science.aal2541 PMID: 28104890.

88. Leicher H, Schade S, Huebbers JW, Munzert-Eberlein KS, Haljiti G, Ludwig C, et al. Endoge-nous RALF peptide function is required for powdery mildew host colonization. biorxiv. 2025. doi:10.1101/2025.01.30.635691

89. Kessler SA, Shimosato-Asano H, Keinath NF, Wuest SE, Ingram G, Panstruga R, et al. Con-served molecular components for pollen tube reception and fungal invasion. Science. 2010; 330(6006): 968–71. doi:10.1126/science.1195211 PMID: 21071669.

90. Torres-Vera R, García JM, Pozo MJ, López-Ráez JA. Do strigolactones contribute to plant de-fence? Mol Plant Pathol. 2014; 15(2): 211–6. doi:10.1111/mpp.12074 PMID: 24112811.

91. Mashiguchi K, Morita R, Tanaka K, Kodama K, Kameoka H, Kyozuka J, et al. Activation of strigolactone biosynthesis by the DWARF14-LIKE/KARRIKIN-INSENSITIVE2 pathway in mycorrhizal angiosperms, but not in Arabidopsis, a non-mycorrhizal plant. Plant Cell Physiol. 2023; 64(9): 1066–78. doi:10.1093/pcp/pcad079 PMID: 37494415.

92. Kodama K, Rich MK, Yoda A, Shimazaki S, Xie X, Akiyama K, et al. An ancestral function of strigolactones as symbiotic rhizosphere signals. Nat Commun. 2022; 13(1): 3974. doi:10.1038/s41467-022-31708-3 PMID: 35803942.

93. Amanda D, Frey FP, Neumann U, Przybyl M, Šimura J, Zhang Y, et al. Auxin boosts energy generation pathways to fuel pollen maturation in barley. Curr Biol. 2022; 32(8): 1798–1811.e8. doi:10.1016/j.cub.2022.02.073 PMID: 35316655.

94. Soneson C, Love MI, Robinson MD. Differential analyses for RNA-seq: transcript-level estimates improve gene-level inferences. F1000Res. 2015; 4:1521. doi:10.12688/f1000research.7563.1

95. Ried MK, Banhara A, Hwu FY, Binder A, Gust AA, Höfle C, et al. A set of Arabidopsis genes involved in the accommodation of the downy mildew pathogen *Hyaloperonospora arabidopsidis*. PLoS Pathog. 2019; 15(7): e1007747. doi:10.1371/journal.ppat.1007747 PMID: 31299058.

96. Medina-Puche L, Tan H, Dogra V, Wu M, Rosas-Diaz T, Wang L, et al. A defense pathway linking plasma membrane and chloroplasts and co-opted by pathogens. Cell. 2020; 182(5):1109–1124.e25. doi:10.1016/j.cell.2020.07.020 PMID: 32841601.

97. Briddon RW, Watts J, Markham PG, Stanley’ J. The coat protein of beet curly top virus is essential for infectivity. Virology. 1989; 172(2): 628–33. doi:10.1016/0042-6822(89)90205-5 PMID: 2800340.

98. Zhang L, Hua C, Pruitt RN, Qin S, Wang L, Albert I, et al. Distinct immune sensor systems for fungal endopolygalacturonases in closely related *Brassicaceae*. Nat Plants. 2021; 7(9):1254–63. doi:10.1038/s41477-021-00982-2 PMID: 34326531.

99. Murray MG, Thompson WF. Rapid isolation of high molecular weight plant ONA. Nucleic Acids Res. 1980; 8(19): 4321–6. doi:10.1093/nar/8.19.4321 PMID: 7433111.

100. Bennett T, Liang Y, Seale M, Ward S, Müller D, Leyser O. Strigolactone regulates shoot development through a core signalling pathway. Biol Open. 2016; 5(12): 1806–20. doi:10.1242/bio.021402 PMID: 27793831.

101. Stirnberg P, van de Sande K, Leyser HMO. MAX1 and MAX2 control shoot lateral branching in Arabidopsis. Development. 2002; 129(5): 1131–41. doi:10.1242/dev.129.5.1131 PMID: 11874909.

